# Guiding ATR and PARP inhibitor combinations with chemogenomic screens

**DOI:** 10.1101/2021.12.13.472393

**Authors:** Michal Zimmermann, Cynthia Bernier, Beatrice Kaiser, Sara Fournier, Li Li, Jessica Desjardins, Alexander Skeldon, Victoria Rimkunas, Artur Veloso, Jordan T. F. Young, Anne Roulston, Michael Zinda

## Abstract

Combinations of inhibitors of Ataxia Telangiectasia- and Rad3-related kinase (ATRi) and poly(ADP-ribose) polymerases (PARPi) synergistically kill tumor cells through modulation of complementary DNA repair pathways, but their tolerability is limited by hematological toxicities. To address this we performed a genome-wide CRISPR/Cas9 screen to identify genetic alterations that hypersensitize cells to a combination of the ATRi RP-3500 with PARPi, including deficiency in RNase H2, *RAD51* paralog mutations or the Alternative Lengthening of Telomeres telomere maintenance mechanism. We show that RP-3500 and PARPi combinations kill cells carrying these genetic alterations at doses sub-therapeutic as single agents. We also demonstrate the mechanism of combination hypersensitivity in RNase H2-deficient cells, where we observe an irreversible replication catastrophe, allowing us to design a highly efficacious and tolerable *in vivo* dosing schedule. Altogether, we present a comprehensive dataset to inform development of ATRi and PARPi combinations and an experimental framework applicable to other drug combination strategies.

## INTRODUCTION

Rational combinations of cytotoxic chemotherapeutics have been long used in cancer treatment as they can synergistically enhance anti-tumor activity compared to single agent regimens and may delay or overcome the emergence of drug resistance (Frei et al., 1965). The recent clinical development of cancer therapies based on synthetic lethal targeting of the DNA damage response (DDR) pathways provides an exciting opportunity for the design of novel drug combinations, since most types of DNA damage can be repaired by parallel pathways and complementary mechanisms of action. DDR targeting compounds are thus prime candidates for eliciting synergistic activity when used in combination (Brown et al., 2017; Pilié et al., 2019; Setton et al., 2021). However, drug combination synergism is often accompanied by increased systemic toxicity, and strategies must be developed to improve tolerability, for example by rational patient selection and/or careful dose-scheduling (Brown et al., 2017; Fang et al., 2019; Pilié et al., 2019).

The first application of synthetic lethality in cancer therapy was the use of poly(ADP-ribose) polymerase inhibitors (PARPi) against *BRCA1/2*-deficient tumors (Bryant et al., 2005; Farmer et al., 2005). PARPi monotherapy has been approved in several clinical settings since the initial discovery of its synthetic lethal potential (Hussain et al., 2020; Pujade-Lauraine et al., 2017; Robson et al., 2017) and multiple PARPi combinations are currently under investigation (Brown et al., 2017; Pilié et al., 2019). Another class of compounds targeting the DDR based on synthetic lethality are inhibitors of Ataxia Telangiectasia- and Rad3-related (ATR) kinase, a key regulator of the replication stress response (Bradbury et al., 2020; Lecona and Fernandez-Capetillo, 2018). Multiple ATR inhibitors (ATRi) are in clinical development as monotherapies and in combination with PARPi, including RP-3500, a novel, highly potent, selective, and orally bioavailable ATRi that has shown robust preclinical efficacy in models deficient in *BRCA1/2* or the Ataxia Telangiectasia-Mutated kinase gene (*ATM*) (Roulston et al., 2021, manuscript in press).

Combining PARPi and ATRi is based on a strong mechanistic rationale, as PARPi create DNA damage that engages the ATR pathway to facilitate cell survival. After activation by replication-associated single-stranded DNA (ssDNA) lesions, ATR coordinates the DDR in multiple ways: First, ATR activates a cell cycle checkpoint to prevent mitosis before DNA damage is repaired. Second, ATR blocks firing of dormant replication origins to avoid further propagation of replication stress. Lastly, ATR mediates stabilization and repair/restart of stalled replication forks by activating fork remodeling enzymes and DNA repair by homologous recombination (reviewed by Saldivar et al., 2017). PARPi create S-phase DNA damage by preventing the release of PARP enzymes from their DNA-bound state, a phenomenon known as ‘PARP trapping’ (Murai et al., 2012), as well as by other mechanisms, e.g., preventing repair of ssDNA replication gaps (Cong et al., 2021). In *BRCA1/2-*deficient cells, ATRi prevent cellular recovery from PARPi treatment and cause rapid cell death by premature mitotic entry with unrepaired DNA damage and inflammatory signaling (Bryant et al., 2005; Farmer et al., 2005; Kim et al., 2017; Schoonen et al., 2019). Consequently, combinations of ATRi and PARPi show synergistic cytotoxicity in cell viability assays and enhanced *in vivo* efficacy over each single agent in pre-clinical *BRCA1/2*- or *ATM*-deficient tumor models (Kim et al., 2020, 2017; Lloyd et al., 2020). Furthermore, in a Phase I clinical study, a combination of the ATRi AZD6738 with the PARPi olaparib showed signals of activity in *BRCA1/2*-mutated tumors (Yap et al., 2016). ATRi have also been shown to overcome several mechanisms of acquired PARPi resistance in preclinical models (Kim et al., 2020; Murai et al., 2016; Yazinski et al., 2017), suggesting a potential application in a PARPi-resistant tumor setting.

Despite the proven synergy, taking full advantage of ATRi/PARPi combinations has been hindered by their limited tolerability due to hematological toxicities (Fang et al., 2019; Yap et al., 2016). Pre-clinical proof-of-concept studies showed that this problem could be overcome by the identification of tumors carrying hypersensitizing alterations, such as *BRCA1/2* or *ATM*-deficiency, which would allow the use of lower doses while maintaining efficacy (Kim et al., 2017; Lloyd et al., 2020). However, a comprehensive genome-wide map of genetic alterations that cause hypersensitivity to ATRi/PARPi combinations is not available. Another approach to mitigate the systemic toxicity is the design of novel dose schedules to minimize the effect of the combination on normal dividing tissues (Fang et al., 2019). Successful clinical development of ATRi/PARPi combinations will likely require application of these two approaches in a concerted fashion.

In recent years, CRISPR/Cas9 chemogenomic screening has proven useful in mapping cellular responses to many compounds used in cancer therapy, including PARPi and ATRi (Hustedt et al., 2019; Olivieri et al., 2020; Wang et al., 2019; Zimmermann et al., 2018). In case of ATRi, a ‘consensus’ set of sensitizing alterations has been proposed by compiling multiple parallel CRISPR screening datasets (Hustedt et al., 2019). Here we employ this technology to chart the cellular factors mediating the response to a ATRi/PARPi combination and identify cancer-relevant genetic alterations that cause profound sensitization. Furthermore, we describe the mechanism by which cells carrying one of these alterations, loss of the RNase H2 enzyme, are affected by the combination and use this information to design an optimized, efficacious *in vivo* dosing schedule that is well tolerated in mouse models. Altogether, we present an experimental framework that can be adapted to many drug combination strategies and show that this framework generates novel insights into the use of ATRi/PARPi combinations that can inform clinical development.

## RESULTS

### A chemogenomic screen for genes underlying the cellular response to single agent and combination treatment with ATR and PARP inhibitors

To map the cellular response to RP-3500 alone or in combination with PARPi we performed a CRISPR chemogenomic screen in Cas9-expressing RPE1-hTERT *TP53* knock out (KO) cells using the TKOv3 library (Hart et al., 2017; Olivieri and Durocher, 2021). We treated cells with dimethyl sulfoxide (DMSO), the ATRi RP-3500, the PARPi niraparib or both in combination (**Figure 1A**). We analyzed data using the CRISPRCountAnalysis (CCA) algorithm (Adam et al., 2021), which stratifies statistically significant drug sensitizer hits into four ‘Jenks classes’, with Jenks class 4 being the strongest hits and Jenks class 1 being the weakest (**Supplementary Table 1**). We identified 122 gene hits in Jenks classes 3 and 4, whose targeting single guide RNA (sgRNA) led to sensitivity to either RP-3500 alone, niraparib alone, or RP- 3500/niraparib. Among these genes, 12 scored as hits in all three treatment arms, whereas several arm-specific hits were identified, including 18 genes specific to the combination arm (**Figure 1B, Supplementary Table 1**). We will hereafter refer to these genes as ‘high-confidence’ hits. When we extended the analysis to all Jenks classes, we obtained a total of 377 hits, of which 46 scored in all arms and 79 were specific to the combination (**Supplementary Figure 1A, Supplementary Table 1**). We will refer to this dataset as the ‘extended’ hit list.

**Figure 1.**
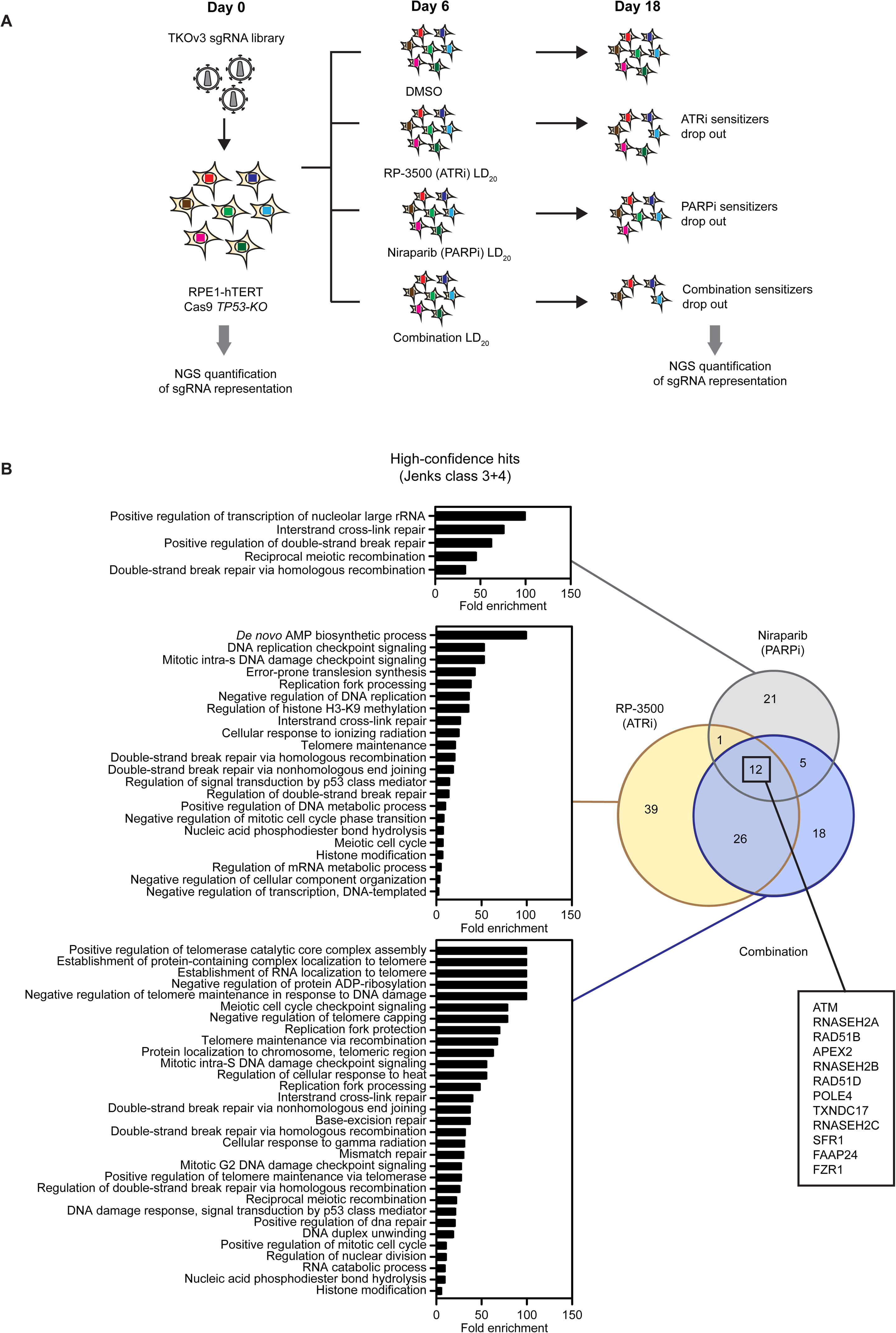
CRISPR screen for sensitizers to RP-3500 + PARPi. **A.** Screen design. Cas9-expressing RPE1 cells were transduced with the TKOv3 library at day 0 and treated with DMSO or LD_20_ doses of each RP-3500, niraparib or combination from days 6 to 18. sgRNA representation at the initial and final timepoints was quantified by NGS. **B.** A Venn diagram showing the number of high-confidence (Jenks class 3+4) hits in the respective treatment arms and GO pathway enrichment in each dataset. Genes scoring in all three arms are highlighted. See also Supplementary Figure 1.

### Robust detection of known DNA repair factors following single agent ATR and PARP inhibitor treatments

The cellular response to single agent ATRi and PARPi has been previously profiled using chemogenomic screening (Bajrami et al., 2014; Hustedt et al., 2019; Olivieri et al., 2020; Wang et al., 2019; Zimmermann et al., 2018) and we could therefore use this information to ‘benchmark’ the ability of our CRISPR-enabled screening pipeline to detect *bona fide* sensitizing gene alterations. We used PANTHER (Mi et al., 2021) to determine in an unbiased way, which Gene Ontology (GO) biological processes were enriched in each arm of our screen (**Figure 1B**). Both single agent ATRi and PARPi arms readily identified homologous recombination (HR) and other genome stability-related processes, confirming the screens’ robustness. Of note, whereas the pathways responding to PARPi treatment were (as expected) centered around HR, several additional ATRi-specific pathways were identified (including translesion synthesis, **Figure 1B**).

Next, we compared our single agent ATRi dataset to the published ‘consensus’ set of ATRi-sensitizing genes from Hustedt et al. (Hustedt et al., 2019) and found 35 overlapping genes (**Supplementary Figure 1B**), including *ATM*, the three RNase H2 subunits *RNASEH2A, B* and *C*, as well as several HR and DNA replication factors (**Supplementary Figure 1B, Supplementary Table 1**). To validate RP-3500 sensitivity after inactivation of selected hits (*ATRIP, RAD17, REV3L, SETD2, CHTF8, FZR1*), we used small interfering RNA (siRNA) pools in two or three human cell lines (RPE1- hTERT Cas9 *TP53-KO*, MCF10A and HeLa). Knockdown of each gene sensitized at least one cell line to RP-3500 to a comparable, or greater, extent than knockdown of *ATM* (**Supplementary Figure 1C**). Collectively, these data confirm that our CRISPR-enabled screening pipeline is robust and able to detect true sensitizers to genotoxic perturbations.

### Identification of genetic vulnerabilities to combined ATR and PARP inhibition

To identify cancer-relevant determinants of sensitivity to ATRi and PARPi combinations we focused on genes scoring in all three treatment arms, as overlapping sensitivity to each single agent should allow achievement of drug synergy at low doses. Among the 12 high-confidence hits were 6 genes with known deleterious alterations in human tumors (Cancer Genome Atlas Research Network et al., 2013): *ATM*, *RNASEH2A, RNASEH2B, RNASEH2C*, and two *RAD51* paralogs, *RAD51B* and *RAD51D* (**Figure 1B, Supplementary Figure 1D, Supplementary Table 1**). A strong vulnerability of *ATM*-deficient tumor cells to combined ATRi and PARPi has been shown previously (Lloyd et al., 2020). RNase H2 is a protein complex composed of three individual polypeptides - the catalytic RNASEH2A subunit and two scaffolding subunits, RNASEH2B and RNASEH2C, in a manner where all three proteins are essential for the stability and activity of the complex (Reijns et al., 2011; Reijns and Jackson, 2014).

RNase H2 plays a key role in the ribonucleotide excision repair pathway, which is responsible for removal of ribonucleotides mis-incorporated into DNA by replicative polymerases (Pizzi et al., 2015; Reijns et al., 2012; Reijns and Jackson, 2014). Biallelic loss of the *RNASEH2B* gene can be found in a subset (∼14%) of chronic lymphocytic leukemias as part of the 13q14 tumor suppressor locus deletion (which includes *RB1* and the *DLEU*/miR-15a/miR-16-1 cluster) (Zimmermann et al., 2018). Possible *RNASEH2B* loss in other tumor types carrying *RB1* homozygous deletions is being investigated (Wang et al., 2019; Zimmermann et al., 2018). *RAD51* paralogs play important roles as chaperones of the RAD51 nucleofilament formation during HR and maintenance of stalled replication forks (reviewed in Bonilla et al., 2020; Rein et al., 2021). Consistent with this function, germline mutations in *RAD51B, RAD51C* and *RAD51D* predispose to breast or ovarian tumors and early PARPi efficacy signals in *RAD51* paralog-mutated tumors are emerging in the clinic (Akbari et al., 2010; Golmard et al., 2013; Loveday et al., 2011; Swisher et al., 2021).

Of note, when mining our extended datasets, we found other factors associated with the same pathways as the 12 high-confidence hits. *NBS1/NBN*, a member of the *MRE11-NBS1-RAD50* (MRN) complex required for ATM activation (Syed and Tainer, 2018) scored as a combination-specific hit (**Supplementary Figure 1D**, **Supplementary Table 1**). Overlapping ATRi and PARPi sensitivities of *MRE11*- deficient cells were shown previously (Fagan-Solis et al., 2020) and we demonstrated efficacy of RP-3500 in a model carrying a *MRE11* alteration (Roulston et al., submitted). Whether the PARPi combination would further enhance efficacy in this model remains to be tested. Unsurprisingly, we also detected numerous HR factors; however, we do highlight that in addition to *RAD51B* and *RAD51D*, we detected all remaining canonical *RAD51* paralogs, including *RAD51C*, *XRCC2* and *XRCC3* in our extended ATRi and ATRi/PARPi datasets (**Supplementary Figure 1D**, **Supplementary Table 1**).

In both *ATM*- and *BRCA*-deficient models, ATRi/PARPi combinations have shown enhanced efficacy over single agent treatments (Kim et al., 2020, 2017; Lloyd et al., 2020). Therefore, next we evaluated whether this would apply also to the additional genetic alterations identified here.

### RNase H2-deficient cells are profoundly sensitive to combined ATR and PARP inhibition

We and others have previously described single agent sensitivity of RNase H2-deficient cells to PARPi and ATRi (Hustedt et al., 2019; Wang et al., 2019; Zimmermann et al., 2018). To validate the combination screen results, we examined whether *RNASEH2B* KO cells had enhanced sensitivity to combined PARPi and ATRi compared to single agent treatment and also their wild type (WT) isogenic counterparts using three independent cellular backgrounds (5637, DLD1 and RPE1-hTERT *TP53-KO*) (Zimmermann et al., 2018), **Figure 2A,B, Supplementary Figure 2A**). As expected, all *RNASEH2B-KO* cells were sensitive to single agent RP-3500 and the PARPi niraparib or talazoparib (**Supplementary Figure 2B-K**). Importantly, combining RP-3500 with PARPi strongly enhanced this cytotoxic effect and doses, which were benign to WT cells and sublethal in *RNASEH2B-KO* cells as single agents, were sufficient to nearly eliminate *RNASEH2B-KO* cells (**Figure 2C-E; Supplementary Figure 3A-D**). We determined that the cytotoxic effect of combined treatment was synergistic by utilizing the SynergyFinder tool (Ianevski et al., 2020; Yadav et al., 2015). Notably, in *RNASEH2B*-*KO* cells concentrations of RP-3500 and PARPi that showed synergy were markedly lower than in WT cells (**Figure 2F-H**; **Supplementary Figure 3E**). Together, these results validate that RNase H2-deficient cells are profoundly sensitive to combinations of PARPi and ATRi.

**Figure 2.**
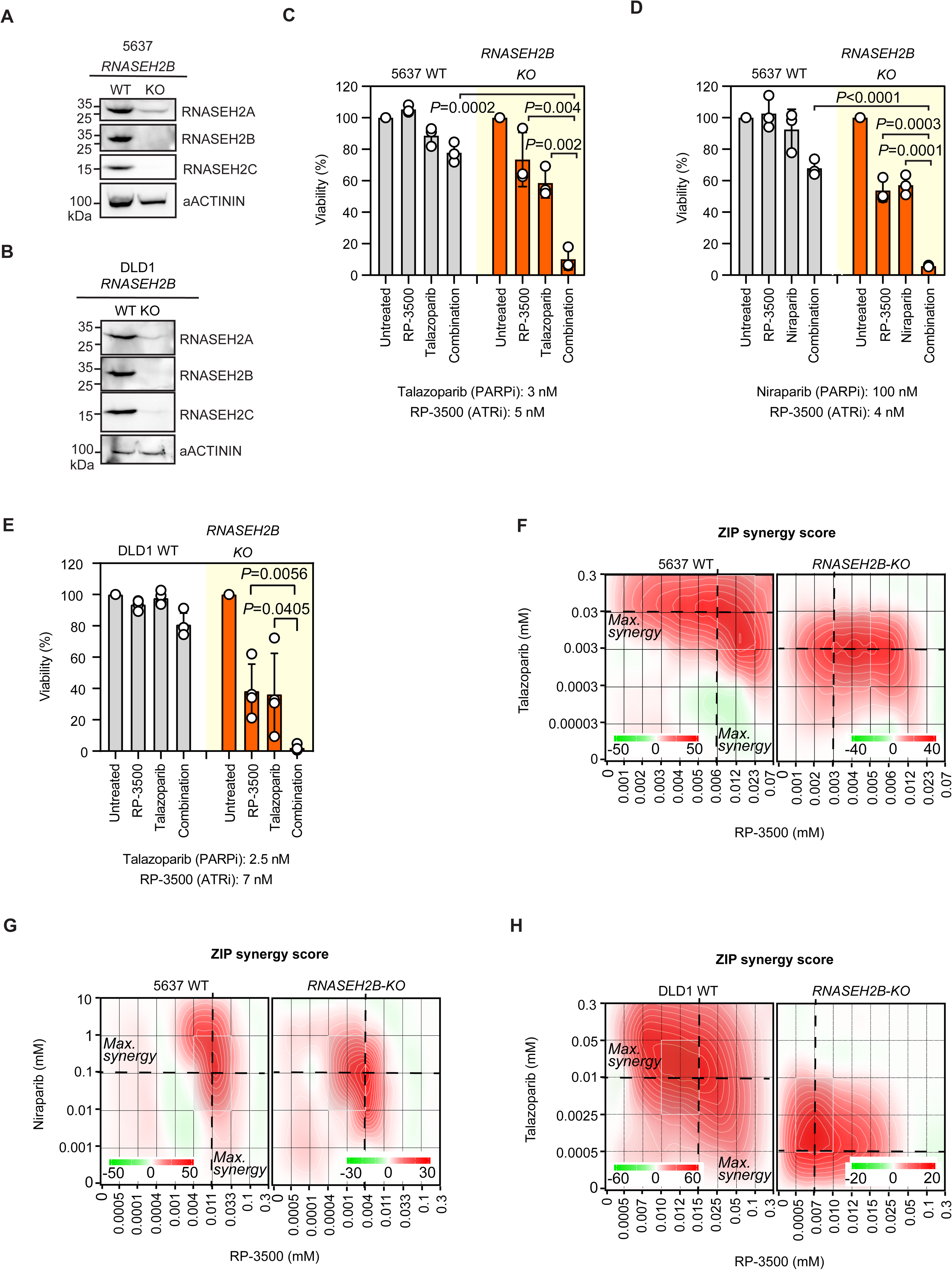
RNase H2-deficient cells are sensitive to ATRi/PARPi. **A,B.** Loss of RNase H2 subunits in *RNASEH2B-KO* cells. Representative (of ≥ independent biological replicates) immunoblots of 5637 (**A**) or DLD1 (**B**) WT and *RNASEH2B-KO* cells. Whole cell lysates were processed for immunoblotting with indicated antibodies. αACTININ, loading control. **C-E.** Viability of 5637 or DLD1 cells of indicated genotypes treated with either RP-3500 (ATRi), PARPi talazoparib or niraparib, or a RP-3500+PARPi combination; values are normalized to untreated controls. Doses of compounds are shown below panels. Circles are values from three independent biological replicates; Bars indicate mean ±SD. *P* values calculated with a two-tailed unpaired Student’s t-test. **F-H.** Zero interaction potency (ZIP) synergy scores at various dose combinations of RP- 3500 and PARPi in cell lines of indicated genotypes. Score ≥10 (red) represents synergy, ≤-10(green) antagonism. Dashed lines mark doses that show maximal synergy. Values obtained by analyzing mean data from three independent biological replicates with SynergyFinder. See also Supplementary Figure 2 and 3.

### Distinct response to PARPi and ATRi in cells lacking RAD51 paralogs

*RAD51* paralogs are required for HR and it is not surprising that they were hits in our screen. However, these proteins have additional and less explored roles in genome stability maintenance, including replication fork protection and DNA damage tolerance, and contributions of the different *RAD51* paralogs to these non-HR functions remain poorly understood (Rein et al., 2021). Interestingly, loss of individual *RAD51* paralogs has been shown to differentially impact PARPi sensitivity *in vitro,* whereby *RAD51B* deficiency displays a milder phenotype than loss of other paralogs (Garcin et al., 2019). As *RAD51B* was one of the strongest ATRi and ATRi/PARPi sensitizers in our screen, we hypothesized that the response of *RAD51* paralogs to ATRi or ATRi/PARPi may differ from that of single agent PARPi. To test this, we depleted *RAD51B, RAD51C or RAD51D* from RPE1-hTERT Cas9 *TP53-KO* cells with lentiviral sgRNAs and confirmed that loss of *RAD51C* and *RAD51D* sensitized more strongly to the PARPi olaparib than *RAD51B,* despite *RAD51B* sgRNAs displaying high cutting efficiency as judged by Inference of CRISPR Edits (ICE) analysis (**Supplementary Figure 4A**; Note that *sgRAD51C-2* that causes lower olaparib sensitivity also induces a lower number of out-of-frame indels; Hsiau et al., 2018). At the same time, the sensitivity to single agent RP- 3500 was comparable between all three paralogs (**Figure 3A-D**).

**Figure 3.**
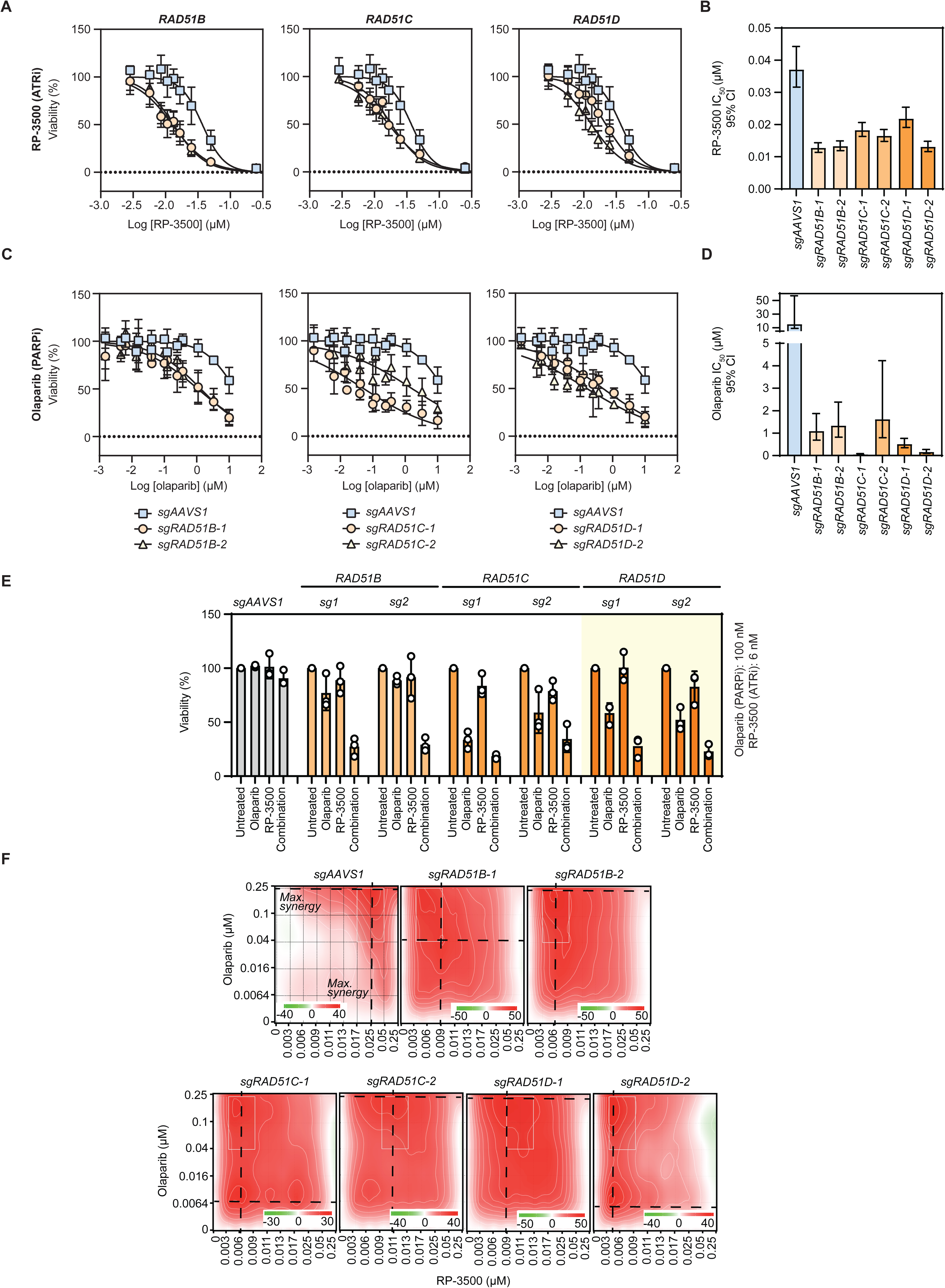
Differential requirement of *RAD51* paralogs in response to PARPi and ATRi. **A-D.** Single agent ATRi (RP-3500, **A,B**) and PARPi (olaparib, **C,D**) sensitivity in cells deficient for *RAD51* paralogs. **A,C.** Dose-response curves of Cas9-expressing RPE1 cells transduced with indicated sgRNAs. Data points are mean of ≥3 independent biological replicates ±SD, normalized to untreated controls. Solid lines show a non- linear least square-fit to a four-parameter dose-response model. **B,D.** RP-3500 and olaparib IC_50_ values in cells transduced with the indicated sgRNAs. Values obtained from non-linear least square fitting of data from three independent biological replicates as shown in **A** and **C**. Error bars represent a 95% confidence interval (CI). **E.** Combination sensitivity. Viability of sgRNA-transduced RPE1 cells after indicated treatments; values are normalized to untreated controls. Doses of compounds are shown below panels. Circles are values from three independent biological replicates; Bars indicate mean ±SD. **F.** ZIP synergy scores at various dose combinations of RP-3500 and olaparib in cells transduced with indicated sgRNAs. Score ≥10 (red) represents synergy, ≤-10 (green) antagonism. Dashed lines mark doses showing maximal synergy. Values obtained by analyzing mean data from three independent biological replicates with SynergyFinder. See also Supplementary Figure 4.

Furthermore, we validated that loss of *RAD51B, RAD51C* and *RAD51D* equally sensitized to a combination of RP-3500 and the PARPi olaparib **(Figure 3E; Supplementary Figure 4B)**. Doses of RP-3500/olaparib that were near indolent in control (AAVS1-targeting) sgRNA-transduced cells substantially decreased viability of cells transduced with sgRNAs targeting *RAD51B, RAD51C* or *RAD51D*, which was accompanied by strong synergy between RP-3500 and olaparib occurring at lower doses than in *sgAAVS1*-transduced cells (**Figure 3E,F; Supplementary Figure 4B**). These data suggest that ATRi or ATRi/PARPi combinations may be universally effective against *RAD51* paralog-mutated tumor cells, unlike single agent PARPi. In contrast, preclinical data suggest that *RAD51B* alterations may reduce PARPi sensitivity compared with alterations in *RAD51C* and *RAD51D* (Garcin et al., 2019), and the use of ATRi or ATRi/PARPi combinations may overcome this issue.

### Tumor cells utilizing alternative lengthening of telomeres are hypersensitive to ATRi/PARPi combinations

In search of other cancer-relevant genetic alterations sensitive to ATRi/PARPi combinations, we noticed that the hits in our ATRi/PARPi screen arm were enriched for several telomere-related GO processes (**Figure 1B**). Closer examination of the combination dataset revealed the presence of several genes implicated in DNA repair reactions at telomeres in cells utilizing the Alternative Lengthening of Telomeres (ALT) pathway (Zhang and Zou, 2020). *FAAP24*, a member of the FANCM DNA translocase complex, was a hit in the high-confidence combination dataset and *FANCM* itself scored in the extended list (**Supplementary Figure 5A**, **Supplementary Table 1**). Moreover, the replication fork remodeling enzyme *SMARCAL1* was a strong hit in the ATRi and ATRi/PARPi screens (**Supplementary Figure 5A**, **Supplementary Table 1**). Both the *FANCM* complex and *SMARCAL1* have been shown to play key roles at ALT telomeres and are known to be regulated by ATR signaling (Bansbach et al., 2009; Lu et al., 2019; Poole et al., 2015; Silva et al., 2019; Singh et al., 2013). Of note, *ATRX*, a chromatin remodeler whose inactivation is strongly associated with ALT (Zhang and Zou, 2020) scored as a combination-specific hit (**Supplementary Table 1**), albeit we were unable to confidently validate this gene using isogenic cell line pairs (data not shown). Nevertheless, due to the presence of multiple ALT-associated factors in our dataset and as ALT has been previously linked to single agent ATRi and PARPi sensitivity (Flynn et al., 2015; Garbarino et al., 2021), we tested whether ALT-positive cancer cells can be targeted with ATRi/PARPi combinations.

We assembled a panel of five ALT-positive (ALT+; U2OS, SAOS2, HS729T, CAL72 and TM31) and five telomerase-positive (TEL+; HT1080, SJSA1, HS683, HOS and HELA) cancer cell lines, and confirmed ALT-status using the recently developed ssTeloC native fluorescence in-situ hybridization (FISH) assay that reflects the level of extrachromosomal C-circles, a hallmark of ALT (**Figure 4A,B**; (Loe et al., 2020)). We also included three non-cancerous telomerase-positive cell lines (RPE1-hTERT, COL-hTERT and MCF10A) and performed cell viability assays with RP-3500, talazoparib, or the combination. In agreement with previous results (Flynn et al., 2015) we observed that ALT+ cell lines were on average more sensitive to ATR inhibition with single agent RP-3500 than the TEL+ cell lines (**Supplementary Figure 5B**). The addition of talazoparib exacerbated the cytotoxic effect of RP-3500, leading to near full loss of viability in ALT+ cell lines at doses that induced a considerably milder effect in TEL+ cancer cell lines and normal immortalized cell lines (**Figure 4C**). One exception was the SAOS2 cell line, which was more resistant to ATRi/PARPi than other ALT+ cell lines as well as TEL+ cells (**Figure 4C**). The reason for this discrepancy is unknown, but it may point to redundancies or heterogeneity among ALT-related mechanisms (Verma et al., 2019; Zhang et al., 2019)

**Figure 4.**
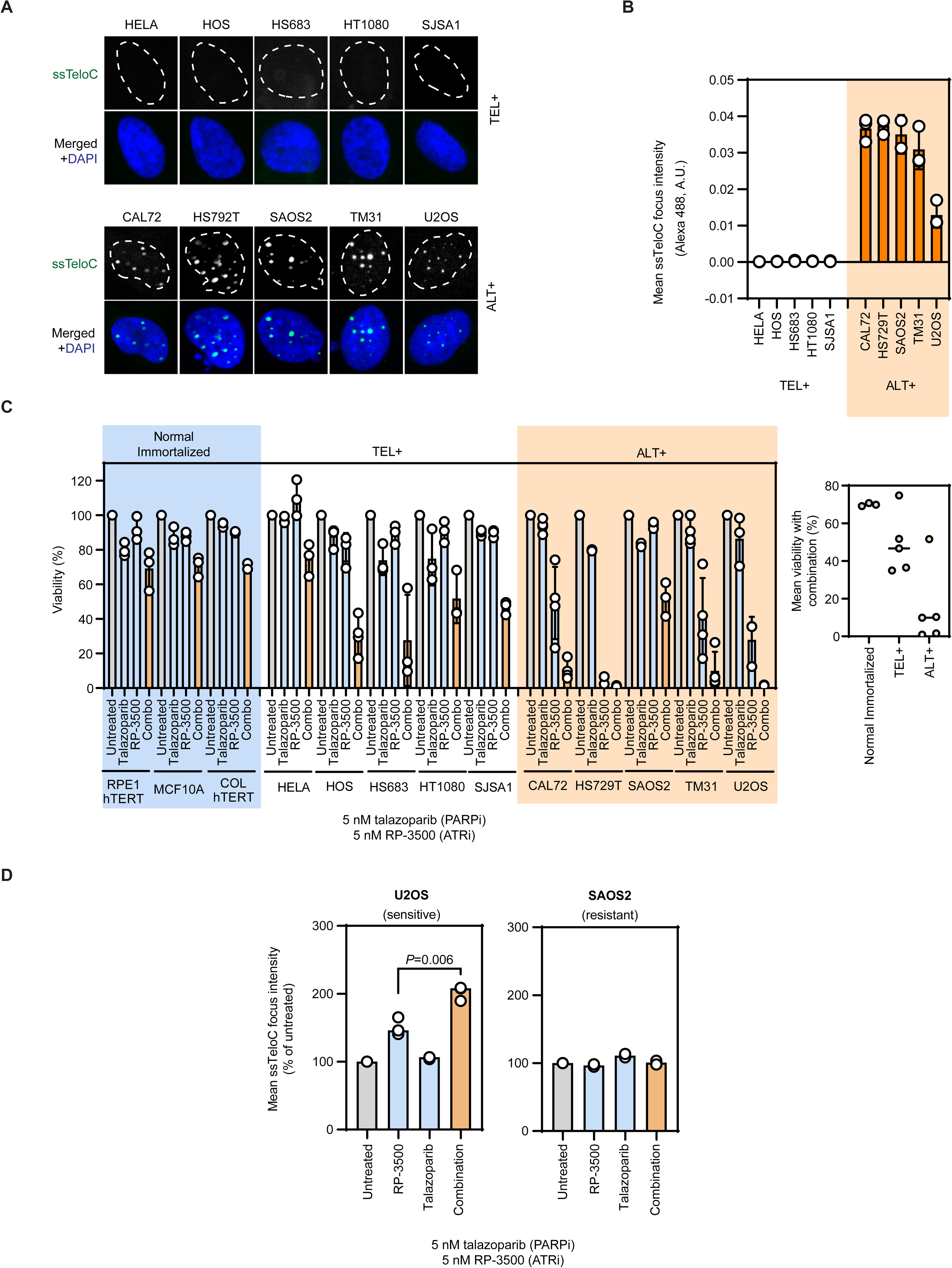
ALT+ cells are sensitive to RP-3500+PARPi. **A.** Representative ssTeloC micrographs showing ALT activity status in the indicated cell lines. ssTeloC foci (green), Alexa Fluor 488; DAPI (blue), nuclear counterstain. **B.** Quantification of mean ssTeloC focus fluorescence intensity in the indicated cell lines. Circles represent mean focus intensity per replicate (N=3 biological replicates, except SAOS2, N=2). Bars show mean ±SD. **C.** Left: Quantification of viability (CellTiter Glo) of the indicated cell lines after treatment with 5 nM RP-3500 (ATRi), 5 nM talazoparib (PARPi) or the combination. Data normalized to untreated cells. TEL+ = telomerase-positive, ALT+ = ALT-positive. Circles represent data from individual biological replicates (N ≥3), bars indicate mean ± SD. Right: Summary view of mean (of three or more biological replicates) combination-treated cell viability in the indicated cohorts. Each data point represents a cell line. **D.** ATRi/PARPi combination elevates ALT activity in a sensitive cell line. Quantification of mean ssTeloC focus fluorescence intensity in a ATRi/PARPi sensitive (U2OS, left) and resistant (SAOS2, right) ALT+ cell line. Circles represent mean focus intensity per replicate (N=3 biological replicates). Bars show mean ±SD. *P* value calculated with a two-tailed unpaired Student’s t-test. See also Supplementary Figure 5.

To exclude the possibility that the ALT+ cell lines in our panel were sensitive to RP-3500+PARPi for reasons unrelated to the ALT phenotype, we evaluated whether ATRi/PARPi combinations affect ALT activity directly. Consistent with that possibility, we observed an increase in ALT activity as measured by ssTeloC in the U2OS ALT+ cell line upon RP-3500 treatment and a further synergistic increase in combination with talazoparib (**Figure 4D**). Interestingly, this effect was not observed in SAOS2 cells, which are resistant to the combination (**Figure 4D**). These results suggest that the sensitivity to the combined ATRi/PARPi correlates with its impact on ALT activity. Excessive ALT activity has been shown to be detrimental due to overwhelming levels of telomere replication stress, as has recently been shown upon inactivation of *FANCM* (Lu et al., 2019; Silva et al., 2019).

### A mechanism of ATRi/PARPi sensitivity in RNase H2-deficient cells

Having identified multiple genetic alterations sensitizing to ATRi/PARPi combination, we next sought to design an optimal dosing schedule for *in vivo* use of RP-3500/PARPi. To achieve this, we first developed a deeper understanding of the mechanism by which the combination affects sensitive genetic backgrounds. We focused on RNase H2 due to the profound sensitivity to the combination in cells mutated for RNase H2-encoding genes. Furthermore, it was unclear how RNase H2-deficient cells are affected by ATRi/PARPi combination in comparison to other alterations such as *ATM* or *BRCA1/2* deficiency (Kim et al., 2017; Lloyd et al., 2020; Schoonen et al., 2019).

We initially monitored cell cycle progression and DNA damage accumulation after PARPi, ATRi and ATRi/PARPi treatment using high-content microscopy in 5637 WT and *RNASEH2B-KO* cells with γ-H2AX as a marker of DNA damage, 4,6-diamidino-2-phenylindole (DAPI) as a measure of DNA content and 5-ethynyl-2′-deoxyuridine (EdU) to label cells replicating their DNA (**Supplementary Figure 6A**). The RP- 3500+niraparib combination led to marked DNA damage accumulation in *RNASEH2B- KO* cells characterized by the appearance of pan-nuclear γ-H2AX (**Figure 5A,B; Supplementary Figure 6A-C**). A substantial portion of pan-γ-H2AX-positive cells showed >2N DNA content but did not incorporate EdU (**Supplementary Figure 6A,B**), suggesting that these cells entered S-phase but are unable to complete DNA replication. Consistent with defective DNA replication, we observed a marked decrease in overall percentage of EdU-positive (EdU+) cells in the combination-treated *RNASEH2B-KO* cells (**Figure 5C; Supplementary Figure 6**). At the same doses, both compounds showed a much milder effect as single agents in *RNASEH2B-KO* cells and no effect was observed in WT cells (even with the combination; **Figure 5A-C; Supplementary Figure 6**). The DNA damage and replication defects observed in *RNASEH2B-KO* cells treated with the RP-3500/niraparib combination were irreversible, as washout did not lower γ-H2AX levels or restore defective EdU incorporation (**Figure 5A-C; Supplementary Figure 6**).

**Figure 5.**
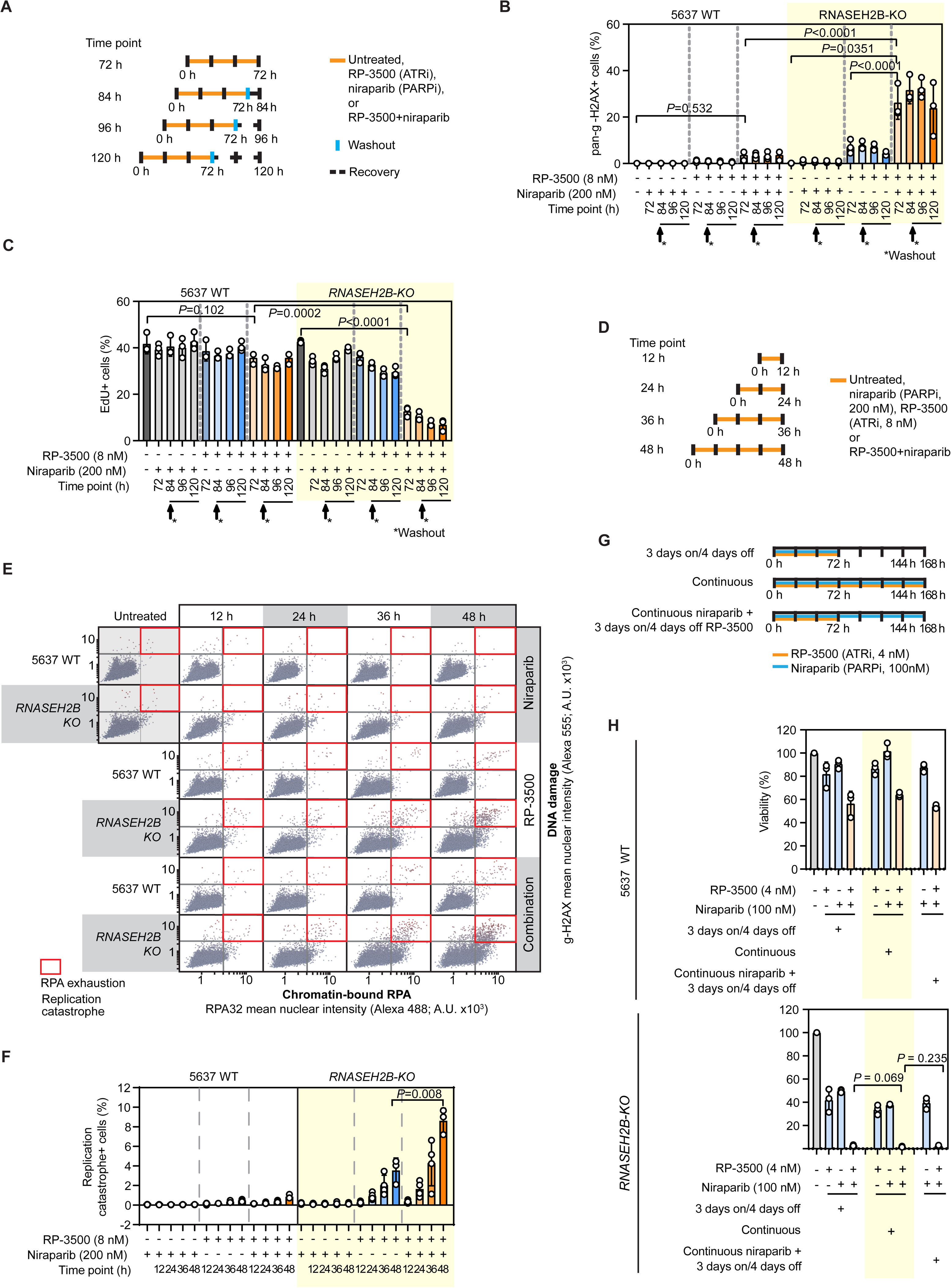
ATRi/PARPi cause irreparable DNA damage and RPA exhaustion in RNase H2-null cells. **A.** Timeline of experiments shown in **B-C**. 5637 WT or *RNASEH2B-KO* cells were treated as indicated for 72 hours followed by immediate processing for immunofluorescence or a washout and a recovery period for indicated time points. **B,C**. Quantification of cells positive (+) for pan-nuclear γ-H2AX (**B**) or EdU (**C**) after indicated treatments and at time points as in **A**. **D.** Timeline of experiments shown in **E,F**. **E.** Quantification of immunofluorescence signals of chromatin-bound RPA32 and γ- H2AX in 5637 WT and *RNASEH2B-KO* cells treated as shown in **D**. Each point represents a single nucleus, solid lines show gates for γ-H2AX+ and RPA32+ cells. Replication catastrophe (RC)+ cells (RPA32+/γ-H2AX+) are outlined with red rectangles. Representative plots of three independent biological replicates. **F.** Percentages of RC+ cells. **G,H**. Short-term ATRi/PARPi treatment is sufficient to kill RNase H2-null cells. **G.** Timeline of the experiment shown in **H.** 5637 WT or *RNASEH2B-KO* cells were treated with RP-3500, niraparib or the combination at the indicated schedules in a 7-day CellTiter Glo viability assay. **H.** Quantification of cell viability in experiments outlined in G. Data in **B,C,F,H** are represented as follows: Circles, values from three independent biological replicates; Bars, mean ±SD. *P* values calculated with a two-tailed unpaired Student’s t-test.

The appearance of pan-nuclear γ-H2AX and an irreversible DNA replication arrest was reminiscent of the previously described phenotype of replication catastrophe (RC) due to replication protein A (RPA) exhaustion (Toledo et al., 2013). To determine whether the ATRi/PARPi combination induces RC in RNase H2-deficient cells, we monitored the levels of chromatin-bound RPA together with γ-H2AX (Toledo et al., 2013). RP-3500+niraparib in both 5637 and RPE1 *RNASEH2B-KO* cells progressively increased the RPA signal accompanied by appearance of high γ-H2AX at later time points, confirming RPA exhaustion and RC (**Figure 5D-F, Supplementary Figure 7A-C**) (Toledo et al., 2013). Consistent with our prior results, we observed a near-total absence of RC in 5637 WT cells following ATRi/PARPi and the effect of the combination was two-fold greater than that of RP-3500 alone in *RNASEH2B-KO* cells (**Figure 5D-F, Supplementary Figure 7A-C**).

RC is an irreversible process leading to cell death (Toledo et al., 2013), suggesting that RNase H2-deficient cells should be killed by the ATRi/PARPi combination within the 48–72 hours needed to induce RC. Indeed, a 72-hour treatment with combined RP-3500 and niraparib followed by removal of the compounds was as lethal to 5637 *RNASEH2B-KO* cells as a continuous, 7-day exposure or continuous niraparib combined with 72 hours of RP-3500 (**Figure 5G,H**). Consistent with these observations, we detected apoptosis marked by caspase-3 cleavage in a subset of *RNASEH2B-KO* cells treated with RP-3500 and niraparib for 72 hours (**Supplementary Figure 7D,E**). Taken together, our results show that a 72-hour treatment with ATRi/PARPi is sufficient to induce RC and cell death in cells lacking RNase H2. Based on these results, we hypothesized that a weekly treatment schedule of intermittent 3 days on/4 days off may be efficacious against RNase H2-deficient tumors *in vivo* and may circumvent the toxicity caused by continuous dosing of ATRi/PARPi (Fang et al., 2019) by limiting the exposure of normal proliferating tissues to the combination.

### *In vivo* activity of ATRi/PARPi combinations

To test whether tumors with RNase H2 loss are sensitive to a low-dose RP-3500/PARPi combination administered at an intermittent schedule *in vivo*, we developed the isogenic DLD1 WT and *RNASEH2B-KO* cells as subcutaneous xenografts in immunocompromised mice. We chose PARPi talazoparib in this study due to its high potency and PARP trapping activity (Murai et al., 2014; Shen et al., 2013) and because it has shown *in vivo* efficacy in an RNase H2-deficient model (Zimmermann et al., 2018). However, the relatively long *in vivo* half-life of talazoparib (de Bono et al., 2017; Stewart et al., 2014) makes its intermittent dosing impractical. Since we saw no difference between 72-hour dosing of both agents and 72-hour dosing of RP-3500 combined with continuous PARPi *in vitro* (**Figure 5G,H**), we evaluated a schedule of continuous talazoparib and RP-3500 dosed intermittently (**Figure 6A**; note that the longer treatment of *RNASEH2B-KO* tumors was to accommodate the slightly slower growth of this model compared to WT). Single agent talazoparib and RP-3500 were dosed at their maximal tolerated doses (MTD), as well as 1/2 MTD of talazoparib or 1/3 MTD of RP-3500 and a combination of the lower doses. The growth of *RNASEH2B-KO* tumors was significantly inhibited by single-agent talazoparib at the MTD, and by RP- 3500 at both MTD and the 1/3 lower dose (∼56%, ∼69% and ∼55% tumor growth inhibition (TGI) respectively, **Figure 6A**) as expected (Wang et al., 2019; Zimmermann et al., 2018). Importantly, the combination of 1/2 MTD talazoparib and 1/3 MTD RP- 3500 led to a marked TGI of ∼83%, which was significantly different from talazoparib at its MTD (**Figure 6A**). At the same time, all agents showed no or only mild activity in RNase H2-proficient tumors (**Figure 6A**). All treatments were well tolerated with <10% mean body weight loss (**Supplementary Figure 8A**).

**Figure 6.**
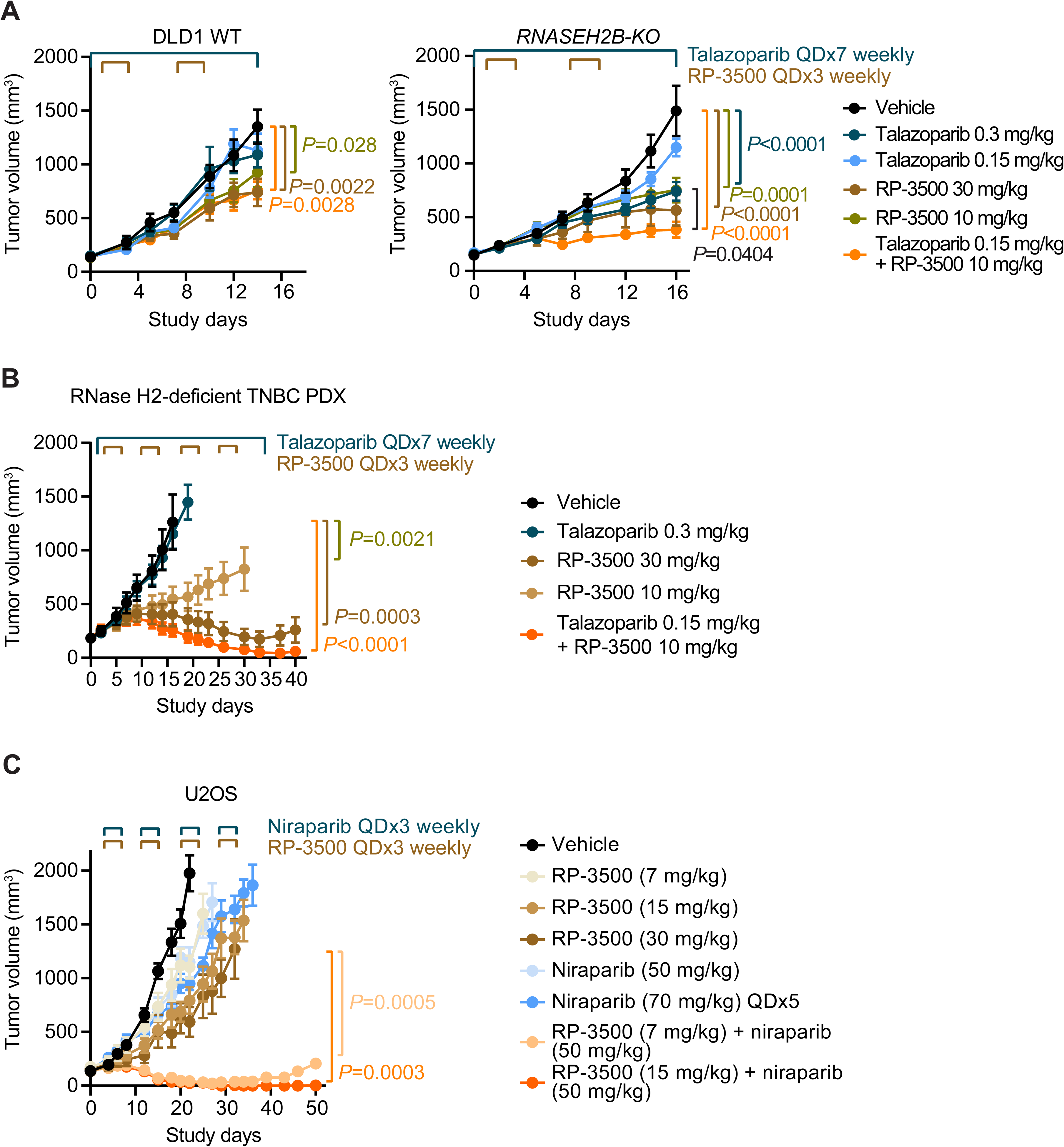
Low dose talazoparib combined with RP-3500 treatment shows efficacy in xenograft models of sensitive genetic backgrounds. **A.** Tumor growth of DLD1 WT and DLD1 *RNASEH2B-KO* cell lines in NOD-SCID mice treated with talazoparib, RP-3500 or both. Talazoparib was administered QDx7 weekly and RP-3500 QDx3 weekly starting on Day 1 of each week. Results are expressed as mean ± SEM, N=8 mice/group and treatment continued for 14 and 16 days for DLD1 WT and DLD1 *RNASEH2B-KO,* respectively. **B.** Tumor growth of a triple negative breast cancer patient-derived PDX in Balb/C nude mice. Talazoparib was administered QDx7 weekly and RP-3500 QDx3 weekly starting on day 2 of every week for 4 weeks. **C.** Tumor growth of U2OS cells in NCG mice treated with niraparib and/or RP-3500 QDx3; niraparib was also given at its MTD of 70 mg/kg QDx5 for comparison. Results are expressed as mean ± SEM, N=8 mice/group. *P* values were determined from tumor volumes by one-way analysis of variance (ANOVA) using the Fisher’s least significant difference (LSD) post-test. See also Supplementary Figure 8.

Next, we tested the efficacy of ATRi/PARPi in a clinically relevant model with loss of RNase H2. Using immunohistochemistry (IHC) with a pan-RNase H2 antibody (Reijns et al., 2012) we identified a patient-derived triple-negative breast cancer tumor xenograft (PDX) without detectable RNase H2 expression (**Supplementary Figure 8B**) and tested the dosing regimens as described above. Single agent talazoparib did not impact growth, but we did observe a dose-dependent tumor growth inhibition with RP- 3500 (**Figure 6B**). Strikingly, the combination of 1/2 MTD talazoparib and 1/3 MTD RP- 3500 led to tumor regression with three of nine mice being tumor free at day 40 as compared to an absence of tumor-free mice in other treatment groups (**Figure 6B**). All treatments were well tolerated with <5% body weight loss (**Supplementary Figure 8C**). Taken together, these data show that RNase H2-deficient tumors are sensitive to ATRi/PARPi combinations *in vivo* and that this sensitivity allows the use of lower doses and intermittent dosing schedules that are well tolerated.

Finally, we tested the *in vivo* activity of a RP-3500/PARPi combination in an ALT+ xenograft tumor model generated from the U2OS cell line. In this case we used the PARPi niraparib, which has a shorter half-life than talazoparib in mice (Jones et al., 2015), and allowed us to directly test a 3 days on/4 days off intermittent dosing schedule. Whereas RP-3500 and niraparib at their MTDs showed only modest efficacy (∼76% and ∼56% TGI, respectively, **Figure 6C**), all combination dosing schedules showed complete tumor regression, with sustained responses for ≥20 days post treatment cessation. Seven of 8 mice were tumor free at day 50 in the 15 mg/kg (1/2 MTD) RP-3500 + 50 mg/kg (∼2/3 MTD) niraparib group, although tumors eventually re- grew (**Figure 6C**). No treatments decreased body weight significantly (**Supplementary Figure 8D**). In summary, these data show that full regression of ALT+ tumors can be achieved with ATRi/PARPi combinations in a mouse model.

## DISCUSSION

PARPi/ATRi combinations hold great promise for cancer therapy. These two classes of agents synergistically enhance each other’s activity against tumor cells and may potentially avoid, delay or overcome therapy resistance (Kim et al., 2020, 2017; Lloyd et al., 2020; Murai et al., 2016; Schoonen et al., 2019; Yap et al., 2016; Yazinski et al., 2017). However, tolerability of the combination at monotherapy MTDs is limited by hematological toxicities (Yap et al., 2016), requiring dose reductions that may impact efficacy. Consequently, there is a clear need to consider alternative strategies to harness the potential of this combination. We contend that these include the development of optimized dosing schedules and/or the identification of sensitivity biomarkers that allow the use of lower doses to improve tolerability, without compromising efficacy.

Here we performed a genome-wide chemogenomic CRISPR/Cas9-based screen to map the cellular response to the novel ATRi RP-3500 as a single agent or in combination with a PARPi. As our screens complement previous datasets (Bajrami et al., 2014; Hustedt et al., 2019; Olivieri et al., 2020; Wang et al., 2019; Zimmermann et al., 2018) we provide it as a public resource, as we believe the data can yield novel biological insights and inform the clinical development of ATRi and ATRi/PARPi combinations. Based on data gleaned here, combined with a proprietary analysis of cancer genomic datasets, we selected 17 ATRi-sensitizing gene mutations (validated here and previously (Hustedt et al., 2019; Wang et al., 2019)) as a basis for patient selection in our first-in-human RP-3500 clinical study (**NCT04497116**).

In our chemogenomic screen we identified and subsequently validated multiple cancer-relevant genetic alterations that profoundly sensitized cells to ATRi/PARPi combinations and may constitute potential clinical biomarkers. Other preclinical studies have identified additional biomarkers for ATRi/PARPi combinations, such as deleterious alterations in *BRCA1/2* (Kim et al., 2020, 2017; Schoonen et al., 2019) or *ATM* (Lloyd et al., 2020), Cyclin E overexpression in breast and ovarian cancer (Kim et al., 2020), *PAX3-FOXO1* fusion in alveolar rhabdomyosarcoma (García et al., 2020), or combined *TP53/RB1* loss in metastatic prostate cancer (Nyquist et al., 2020).

The first cancer-relevant genetic alteration identified here was loss of RNase H2, which expands on previously reported findings of single agent PARPi and ATRi sensitivity (Wang et al., 2019; Zimmermann et al., 2018). We demonstrate that RP- 3500/PARPi combinations synergize and induce cell death in RNase H2-deficient cells at concentrations that are almost benign in cells with functional RNase H2. Furthermore, low-dose RP-3500/PARPi combinations show robust *in vivo* efficacy in RNase H2- deficient xenograft models. Loss of RNase H2 is found in a subset of chronic lymphocytic leukemias (CLL) (Zimmermann et al., 2018); therefore, an important avenue for further study is to test the ATRi/PARPi combinations in preclinical CLL models (Herman and Wiestner, 2016; Kwok et al., 2016; Zimmermann et al., 2018). Loss of *RNASEH2B* was also reported in metastatic prostate cancer, although homozygous deletions of *RNASEH2B* in this context are likely rare (Wang et al., 2019; Zimmermann et al., 2018). It will be crucial to determine if *RNASEH2B* loss can be detected in additional tumor types and whether these tumors can respond to ATRi/PARPi combinations. We have identified a TNBC PDX model with no RNase H2 expression by IHC, suggesting that RNase H2 loss can be detected outside the context of CLL. Of note, while this manuscript was under preparation, Miao et al. reported, in a pre-print, robust efficacy with an ATRi/PARPi combination in RNase H2-deficient prostate cancer models, which was well in agreement with the results of our study (Miao et al., 2021).

Using RNase H2-deficient cells as an example, we showed that the ATRi/PARPi combination can lead to RPA exhaustion and RC. In *BRCA1/2-* or *ATM-*deficient cells, the potentiation of the PARPi effects by ATRi is attributed primarily to unscheduled entry into mitosis due to abrogation of the G2/M phase checkpoint (Kim et al., 2017; Schoonen et al., 2019). Therefore, RNase H2 deficiency is to our knowledge, the first context in which RC contributes to the synergy between ATRi/PARPi. This raises the possibility that PARPi/ATRi combinations can kill cells by multiple (but not necessarily mutually exclusive) mechanisms dependent on the respective genetic context. Importantly, we used our mechanistic findings to design an optimized, 3 days on/4 days off weekly intermittent schedule for *in vivo* dosing of RP-3500/PARPi combinations and demonstrated that this schedule is effective and well tolerated in animal models. We are currently evaluating this and related schedules in Phase I clinical trials (**NCT04497116, NCT04972110**).

Next we showed ATRi/PARPi sensitivity upon inactivation of RAD51 paralogs *RAD51B,C* and *D*. Interestingly, cells lacking *RAD51B* have been shown to be less sensitive to PARPi than cells lacking *RAD51C* and *D*, which has been attributed to a differential requirement of these factors for HR (Garcin et al., 2019). In contrast, we observed that ATRi sensitivity after loss of all three paralogs was comparable, as was the sensitivity to ATRi/PARPi, which cannot be explained by a simple difference in requirement for each paralog for HR. *RAD51* paralogs also play important roles in replication fork maintenance and DNA damage tolerance (Berti et al., 2020; Rein et al., 2021; Saxena et al., 2019; Somyajit et al., 2020), and it is therefore tempting to speculate that the difference in sensitivity to ATRi and PARPi between individual *RAD51* paralogs may be governed by these non-HR functions, although further experimental evidence is needed to support this hypothesis. In a more general sense, our observations highlight the need to establish *bona fide* clinical biomarkers for ATRi. It is apparent from our screen that, despite substantial (and expected) overlap between the ATRi and PARPi treatment arms, some genes and pathways preferentially respond to one treatment over another, and it will be important to establish whether clinical biomarkers for PARPi will be universally applicable to ATRi and *vice versa*.

As a third cancer alteration that sensitizes cells to ATRi/PARPi combinations, we validated ALT. A functional link between ALT and ATRi sensitivity was reported previously, although it remained somewhat controversial (Deeg et al., 2016; Flynn et al., 2015). We identified several ALT-associated factors in our ATRi/PARPi screen and showed that ATRi/PARPi treatment elevates markers of ALT activity, suggesting that the effects of ATRi/PARPi in ALT+ cell lines are due to a direct impact on ALT. In future research it will be important to establish which ATR and PARP targets impact ALT activity; however, ATR-regulated replication fork maintenance factors such as SMARCAL1 or FANCM are obvious candidates (Bansbach et al., 2009; Singh et al., 2013). Interestingly, we showed a strong and durable response of an ALT+ mouse xenograft to the RP-3500/PARPi combination, despite this model showing only moderate responses to each single agent, which highlights the synergistic potential of ATRi/PARPi combinations. It will be important to determine whether other genetic lesions in cancer can create such a profound vulnerability to ATRi/PARPi (and other DDR agent combinations), despite leading to only modest single-agent activity.

In conclusion, we demonstrate that using CRISPR chemogenomic screening combined with mechanistic insights can guide the rational use of drug combinations in preclinical models to minimize toxicity while preserving the efficacy of the combination, and believe that the findings reported here will inform investigation of ATRi/PARPi combination regimens in the clinic.

## AUTHOR CONTRUBUTIONS

M. Zimmermann conceived the study with input from AR and M. Zinda. M. Zimmermann also performed most cell biology experiments, analyzed data and wrote the paper with input from other authors. CB and JD performed the CRISPR screen under supervision from JTFY. BK and AS performed cell biology assays. SF and LL conducted *in vivo* mouse studies with supervision from AR. AV analyzed CRISPR screening and PDX genomic data. VR directed IHC experiments. M. Zinda supervised all activities and helped write the manuscript.

## MATERIALS AND METHODS

### Cell culture and generation of CRISPR KO cell lines

5637, DLD1, RPE1-hTERT, MCF10A, HELA, HOS, HS683, HT1080, SJSA1, HS729T, SAOS2 and U2OS cell lines were purchased from ATCC. CAL72 cells were obtained from DSMZ. COL-hTERT (Immortalized Human Colon Cells) were purchased from ABM. TM31 were obtained from RIKEN BioResource Research Center. 5637, DLD1 and SJSA1 cells were cultured in RPMI-1640 medium (Corning 10-040-CV) supplemented with 10% fetal bovine serum (FBS; VWR 97068-085), 100U/mL penicillin and 100μg/mL streptomycin (Pen/Strep; Corning 30-001-CI). RPE1, TM31, HELA, HT1080, HS729T cells were cultured in Dulbecco’s Modified Eagle Medium (DMEM; Corning 10-014-CV) supplemented with 10% FBS and Pen/Strep. SAOS2 and U2OS cells were cultured in McCoy’s 5A medium (Cytiva SH30200.FS) supplemented with 10% FBS and Pen/Strep. All cell lines were maintained at 37°C and 5% CO_2_.

DLD1 *RNASEH2B-KO* and 5637 *RNASEH2B-KO* cell lines were generated by GenScript USA, Inc., using a 5’ AAGAGAACTTACCTGAACAG 3’ sgRNA target sequence. Briefly, Cas9/sgRNA nucleoprotein complexes were transfected into cells and single clones were isolated by limiting dilution. Clones were screened using polymerase chain reaction (PCR, forward primer 5’ ACCCCTGCTTCTCATCATTCC 3’; reverse primer 5’ TTGCCCGTATTTCTGATGGCT 3’) and TIDE analysis (Brinkman et al., 2014) as well as immunoblotting. RPE1-hTERT Cas9 *TP53-KO* and RPE1-hTERT Cas9 *TP53-KO RNASEH2B-KO* cells were described previously (Zimmermann et al., 2018).

### Chemical compounds

Stock solutions of RP-3500 (synthesized by OmegaChem), olaparib, talazoparib and niraparib (all MedChem Express) were made up from powder in DMSO and kept at -20°C for long-term storage.

### CRISPR/Cas9-enabled chemogenomic screening

CRISPR screens were performed according to a published procedure (Olivieri and Durocher, 2021). On day -3, Cas9-expressing RPE1-hTERT *TP53-KO* cells were infected with the TKOv3 lentiviral library (Hart et al., 2017) at a low multiplicity of infection (MOI; ∼0.6) and transductants were selected with puromycin from day -2 to day 0. On day 0 transduced cell pools were split into two technical replicates and cultured until day 6. On day 6, each replicate was split into four treatment arms – DMSO control, RP-3500, niraparib and a RP-3500+niraparib combination. The following concentrations were used, which resulted in a ∼20% loss of cell viability (an LD_20_ dose) at the screen: 15nM RP-3500 single agent, 1μM niraparib single agent and a combination of 8nM RP-3500 with 150nM niraparib. Cells were cultured in the presence of compounds until day 18. Genomic DNA was isolated from day 0 and day 18 samples, sgRNA genomic integrants were amplified by PCR and sgRNA representation in each sample was quantified by next generation sequencing. Data were analyzed using the CCA algorithm (Adam et al., 2021).

### Lentiviral sgRNA transduction

Individual TKOv3 sgRNAs were cloned into a lentiGuide-NLS-GFP vector (Noordermeer et al., 2018). Lentiviral stocks were prepared by seeding 18 x 10^6^ 293T LentiX cells on a 15cm dish in 20mL media and transfected 24 hours later by adding 2.4mL of transfection mix, which contained OptiMEM media, 18μg of sgRNA plasmid, lentiviral packaging vectors (11.7μg psPAX2 + 6.3μg psPAX2), 90μl Lipofectamine (Thermo Fisher Scientific) and 60μl PLUS Reagent (Thermo Fisher Scientific 11514015). Medium was exchanged ∼16–20 hours after transfection and supplemented with 20mM HEPES. Virus-containing supernatant was collected ∼48 hours post transfection, cleared by centrifugation at 1000 RPM for 5 minutes and stored at -80°C.

1 x 10^5^ RPE1-hTERT Cas9 *TP53-KO* cells were transduced with the virus at MOI ∼1. Virus was removed 24 hours post infection and sgRNA-expressing cells were selected with puromycin-containing media for 48 hours. Transduced cells were seeded for experiments 7 days post infection.

### CellTiter Glo cell viability assays

On day 0, cells were seeded on 96-well plates (Corning, 3903). The following cell numbers and assay lengths were used: 5637 WT: 300 cells/well, 7 days; 5637 *RNASEH2B-KO*: 800 cells/well, 7 days; RPE1-hTERT Cas9 *TP53-KO*: 200 cells/well, 6 days; RPE1-hTERT Cas9 *TP53-KO RNASEH2B-KO*: 200 cells/well, 6 days; DLD1 WT: 300 cells/well, 8 days; DLD1 *RNASEH2B-KO*, 300 cells/well, 9 days.; RPE1-hTERT Cas9 *TP53-KO* infected with lentiviral sgRNAs: 600-800 cells/well, 7 days; HELA: 1000 cells/well, 7 days; HOS: 1000 cells/well, 7 days; HS683: 800 cells/well, 15 days; HT1080: 800 cells/well, 7 days; SJSA1: 800 cells/well, 7 days; CAL72: 1000 cells/well, 9 days; HS729T: 1000 cells/well, 14 days; SAOS2: 800 cells/well, 14 days; TM31: 1000 cells/well, 12 days; U2OS: 800 cells/well, 7days. Compounds were added from DMSO stock solutions on day 1 using a Tecan D300E dispenser (Tecan). Cells were either grown in continuous presence of compounds (medium and compounds were refreshed every 3–4 days) or at various schedules indicated in the respective figures (see results section). Cell viability was measured using the CellTiter Glo assay kit (Promega) according to manufacturer’s instructions. Luminescence was read out either on Envision 2105 (Perkin-Elmer) or FlexStation 3 (Molecular Devices) plate readers. Relative cell viability was calculated by subtracting blank luminescence from each measured value, averaging technical replicates and normalizing to untreated cells. Synergy between RP- 3500 and PARPi was analyzed with the on-line SynergyFinder tool (Ianevski et al., 2020) using the ZIP (Yadav et al., 2015) model (https://synergyfinder.fimm.fi).

### Immunoblotting

Whole cell lysates were prepared by resuspending cell pellets in 2 x Sample Buffer (Novex Tris-Glycine SDS Sample Buffer, ThermoFisher #LC2676, supplemented with 200mM DTT) at a concentration of 1x 10^7^ cells/mL. Lysates were boiled at 95°C for 5 minutes and 15–30μl was separated on Tris-Glycine SDS-PAGE gels (ThermoFisher) followed by Western blotting onto nitrocellulose membranes (ThermoFisher, PB7220) in 1x Novex Tris-Glycine Transfer Buffer (ThermoFisher, LC3675) containing 20% methanol and 0.04% SDS. Membranes were blocked in 5% milk / Tris-buffered saline / 0.1% Tween 20 (5% milk / TBST) and incubated with primary antibodies diluted in 5% milk / TBST either overnight at 4°C or for 2 hours at room temperature (RT). Membranes were washed for 3x 5 minutes with TBST and incubated for 1 hour at RT with horseradish peroxidase-conjugated secondary antibodies (goat anti-rabbit IgG, Jackson ImmunoResearch 111-035-144 or sheep anti-mouse IgG, GE Healthcare NA931V) diluted 1:5000 in 5% milk / TBST. Membranes were washed as above, developed with a SuperSignal West Femto chemiluminescence reagent (ThermoFisher, PI34095) for 2 minutes and scanned on a ChemiDoc imager (Bio-Rad).

### High-Content Fluorescence Microscopy

Cells were grown on CellCarrier Ultra 96-well plates (Perkin Elmer, 6055302) and subjected to various treatments as indicated in the respective figures. If applicable, 10μM EdU was added 30 minutes before fixation to monitor DNA replication. Medium was subsequently removed and cells were rinsed with phosphate buffered saline (PBS). In experiments analyzing the levels of chromatin-bound RPA32 and γ-H2AX (Figure 5D-F, **Supplementary Figure 7**), plates were placed on ice and soluble nuclear proteins were extracted with ice-cold cytoskeleton (CSK) buffer (10mM PIPES pH 7.0, 300mM sucrose, 100mM NaCl, 3mM MgCl_2_) for 15 minutes followed by a rinse with PBS. In all other experiments this CSK pre-extraction step was omitted. Cells were then fixed with 4% paraformaldehyde (PFA) in PBS for 10 minutes at RT and permeabilized with 0.3% Triton X-100/PBS for 30 minutes at RT (permeabilization was omitted if CSK extraction was performed before fixation). Cells were subsequently blocked with PBS/bovine serum albumin/gelatin (PBG: 0.2% cold water fish gelatin, 0.5% bovine serum albumin [BSA] in PBS) for 30 minutes at RT and incubated with primary antibodies diluted in PBG for 2 hours at RT. Afterwards cells were rinsed twice with PBS and incubated with fluorescently labeled secondary antibodies (Alexa Fluor 488 or 555- conjugated goat anti-rabbit IgG or goat anti-mouse IgG, Invitrogen) in PBG for 1 hour at RT. 0.5μg/mL DAPI was included with secondary antibodies as a nuclear counterstain. Cells were finally rinsed 3x with PBS and either immediately imaged or processed for click chemistry to visualize DNA-incorporated EdU. For EdU staining, cells were fixed again with 4% PFA/PBS for 5 minutes at RT and rinsed with PBS. Click chemistry was then performed using a Click-iT EdU Cell Proliferation Kit for Imaging, Alexa 647 (ThermoFisher, C10340), according to the manufacturer’s instructions. Plates were imaged on an Operetta automated microscope (Perkin Elmer) in confocal mode and image analysis was performed in the Harmony software (Perkin Elmer). Examples of gating cells positive for the respective markers is shown in Figure 5E and **Supplementary Figures 6,7**.

### ssTeloC assay

The ssTeloC assay was performed as described previously (Loe et al., 2020). Cells were seeded on CellCarrier Ultra 96-well plates (Perkin Elmer, 6055302) and, if desired, treated with RP-3500, talazoparib, or their combination for 24 hours. Medium was removed, cells were rinsed with PBS and fixed with 2% PFA/PBS for 5 minutes at RT. The plates were then washed with PBS and incubated with RNase A blocking solution (500μg/mL RNase A, 1mg/mL BSA, 3% normal goat serum, 0.1% Triton X-100, 1mM EDTA in PBS) for 1 hour at 37°C. Samples were subsequently dehydrated with a 70%, 95% and 100% ethanol series (5 minutes, RT, each) and allowed to air-dry. Native fluorescence *in situ* hybridization (FISH) was performed by incubating cells with a hybridization mix (70% formamide, 1mg/mL Roche FISH blocking reagent 11096176001, 10mM Tris/HCl pH 7.0) containing an Alexa Fluor 488-labeled TelG PNA probe (PNA BIO F1008) at a 1:500 concentration for 2 hours at RT. Plates were subsequently washed 2x with 70% formamide/PBS for 15 minutes each and 3x 5 minutes with PBS. 0.5μg/mL DAPI was included in the second PBS wash. Plates were imaged on an Operetta automated microscope (Perkin Elmer) in confocal mode and image analysis was performed in the Harmony software (Perkin Elmer).

### Antibodies

The following primary antibodies were used for immunoblotting (IB) and immunofluorescence (IF): Mouse anti-RNASEH2A G-10 (SCBT sc-515475, IB 1:50), rabbit anti-RNASEH2B (Novus NBP2-58962, IB 1:200), rabbit anti-RNASEH2C (Proteintech 16518-1-AP, IB 1:1000), rabbit anti-pCHK1(S345) 133D3 (CST 2348, IB 1:1000), mouse anti-CHK1 2G1D5 (CST 2360, IB 1:200), mouse anti-αACTININ AT6/172 (Millipore Sigma 05-384, IB 1:5000), mouse anti-γ-H2AX JBW301 (Millipore Sigma 05-636, IF 1:2000), rabbit anti-γ-H2AX (CST 2577, IF 1:500), mouse anti-RPA32 (Abcam ab2175, IF 1:1000), rabbit anti-cleaved caspase-3 (CST 9664, IF 1:1000), mouse anti-pH3(S10) (ThermoFisher, MA515220, IF 1:1000).

### Immunohistochemistry

Formalin-fixed, paraffin-embedded PDX tumor samples were obtained from Crown Bio. RNase H2 protein expression was assessed by IHC using a sheep polyclonal anti- RNase H2 antibody (Reijns et al., 2012) (a gift from Andrew Jackson) at 1:300 and rabbit anti sheep IgG (H+L) secondary link antibody (Invitrogen #31240 1:4000) following Leica BondRx standard IHC protocols. PDX models were considered RNase H2-negative if they showed a complete absence of RNase H2 staining.

### Mouse studies

DLD1 and DLD1 *RNASEH2B-KO* cells were implanted at 5 and 10 x 10^6^ cells per mouse, respectively, in PBS into the right flank of female NOD-SCID mice (5–7 weeks old; Charles River). U2OS cells were implanted at 1×10^7^ cells in 50:50 PBS:Matrigel (Corning, # CB35248). When tumors had reached the target size of 100–150mm^3^, mice were randomized into treatment groups (n=7–8). *In vivo* studies involving cell-derived xenografts were performed in a Canadian Council on Animal Care-accredited vivarium with an Institutional Animal Care Committee-approved protocol. *In vivo* studies using PDX were conducted at Crown Biosciences Inc. (Taicang). Fresh tumor tissue from mice bearing established primary human tumors were harvested and cut into small pieces (approximately 2–3mm in diameter). These tumor fragments were inoculated subcutaneously into the upper right dorsal flank of female BALB/c nude mice (5–7 weeks old) for tumor development. When the mean tumor size reached approximately 150 (100–200)mm^3^, mice were randomized into treatment groups (n=6) according to growth rate. The procedures involving the care and use of animals in this study were reviewed and approved by the Institutional Animal Care and Use Committee of CrownBio and were conducted in accordance with the regulations of the Association for Assessment and Accreditation of Laboratory Animal Care.

RP-3500 and niraparib were formulated in 0.5% methylcellulose, 0.02% sodium lauryl sulfate. Talazoparib was formulated in 0.5% carboxymethyl cellulose and combined with RP-3500 before oral administration. Talazoparib was administered once daily (QD) for a maximum of 3 weeks; RP-3500 and niraparib were administered QD for three consecutive days weekly starting on Day 1 or 2 of each week. Tumor volume (TV) was measured using a digital caliper and calculated using the formula 0.52 x Length x Width^2^. Response to treatment was evaluated for TGI. TGI was defined as: %TGI= ([TVvehicle/last – TVvehicle/day0] – [TVtreated/last – TVtreated/day0]) / (TVvehicle/last – TVvehicle/day0) x100. Body weight (BW) was represented as change in BW using the formula: %BW change = (BWlast / BWday0) x 100. BW change was calculated based on individual BW changes relative to Day 0. Statistical significance relative to vehicle control was established by one-way ANOVA followed by Fisher’s LSD test.

## SUPPLEMENTARY FIGURE LEGENDS

**Supplementary Table 1 will be available at publication.**

## ACKNOWLEDGEMENTS

We thank Daniel Durocher and members of the Repare Therapeutics team for critical reading of the manuscript and Daniel Durocher and Andrew Jackson for reagents.

---

**Supplementary Figure 1.**
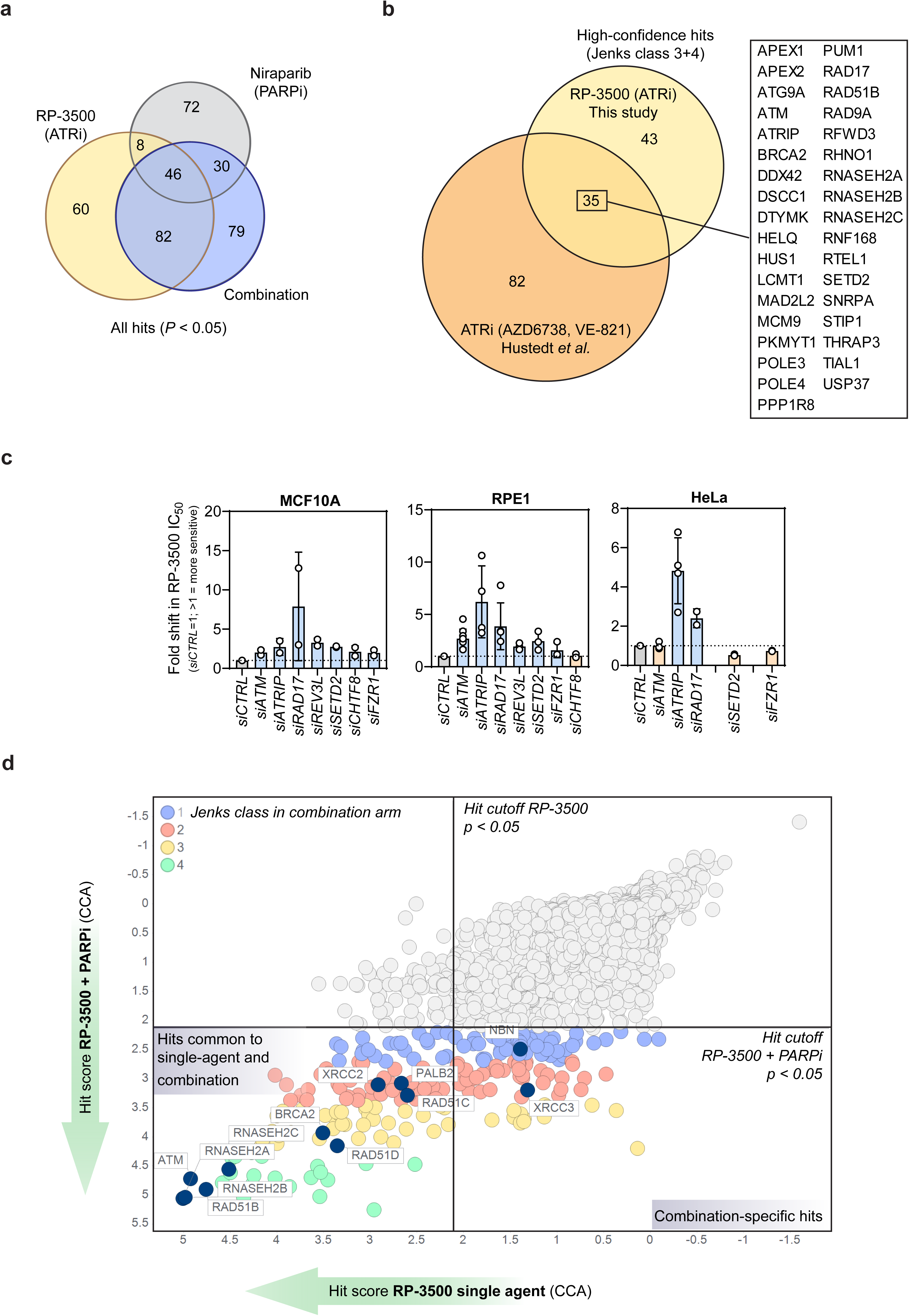
Related to Figure 1. **A.** Venn diagram showing the number of statistically significant hits (*P* <0.05) in the three CRISPR screen treatment arms. **B.** Overlap between the RP-3500 CRISPR screen hits in this study and a published ‘consensus’ set of ATRi sensitizing genes (Hustedt et al., 2019). Common hits are listed on the right. **C.** siRNA validation of selected RP-3500 sensitizing hits. MCF10A, RPE1-hTERT *TP53- KO* and HeLa cells were transfected with indicated siRNA pools and sensitivity to RP- 3500 was determined by 5 day CellTiter Glo assays. Data are represented as a fold-shift in RP-3500 IC_50_ from to non-targeting *siCTRL* pool (circles) from ≥2 independent biological replicates. Bars show mean±SD. **D.** CRISPR screen hits in RP-3500 single agent and RP-3500+PARPi arms as determined by CCA. Each circle represents a gene, straight lines denote hit cutoffs (*P* <0.05). Colors indicate Jenks classes in the RP-3500+PARPi combination arm. Hits discussed in this manuscript are highlighted in dark blue.

**Supplementary Figure 2.**
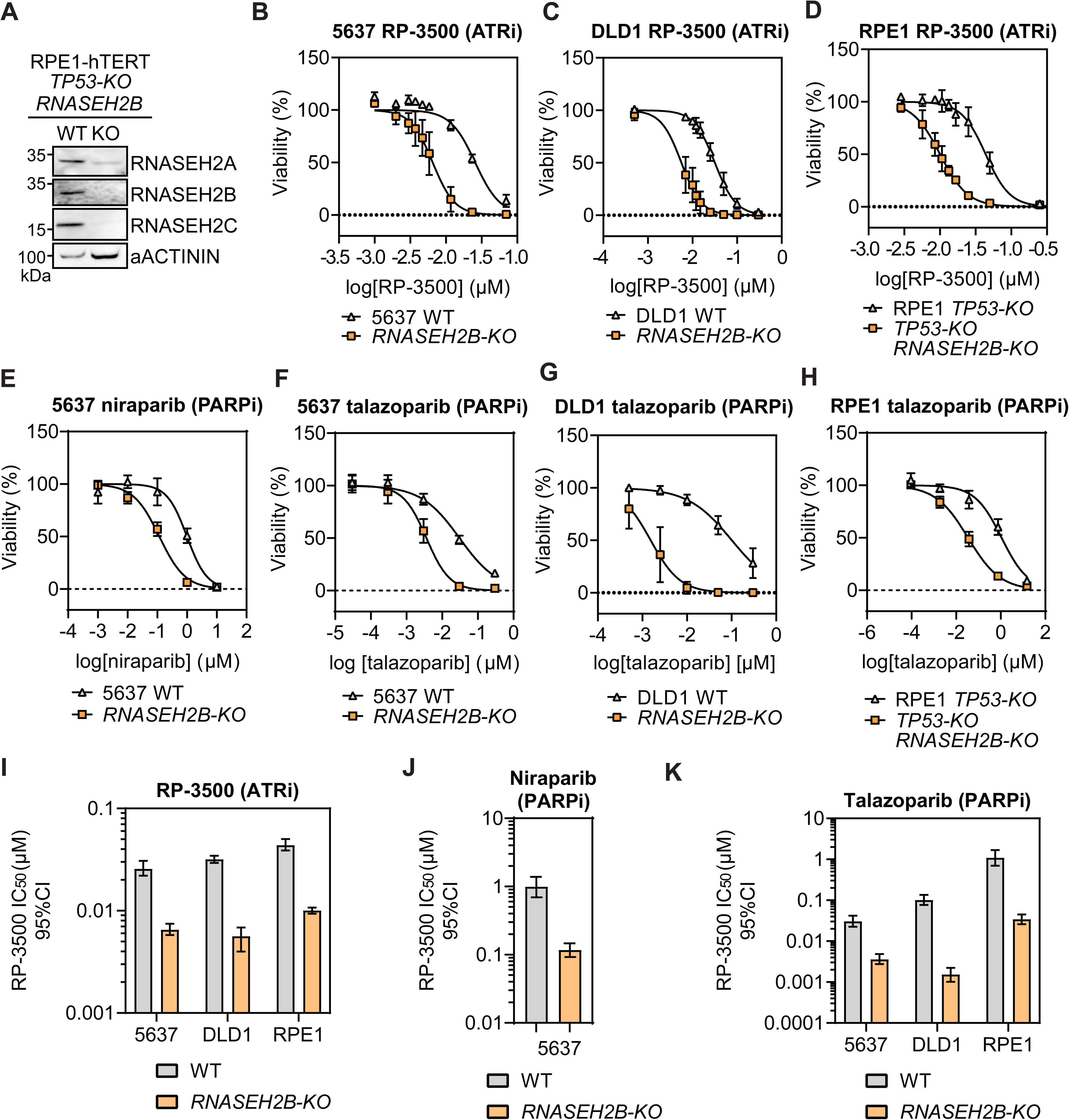
Related to Figure 2. **A.** Loss of RNase H2 subunits in RPE1-hTERT *TP53-KO RNASEH2B-KO* cells. Whole cell lysates of indicated genotypes were processed by immunoblotting with RNASEH2A, B and C antibodies. αACTININ was a loading control. **B-H.** Dose-response curves of 5637, DLD1 and RPE1 cells of indicated genotypes to single agent ATRi (RP-3500, **B-D**) or PARPi (niraparib or talazoparib, **E-H**). Data points are mean of ≥3 independent biological replicates ±SD, normalized to untreated controls. Solid lines show a non-linear least square-fit to a four-parameter dose-response model. **i-k.** RP-3500, niraparib and talazoparib IC_50_ values (shown as bars) for various WT and *RNASEH2B-KO* cell lines from experiments shown in **B-H**. Error bars indicate 95% confidence intervals from non-linear least square fitting.

**Supplementary Figure 3.**
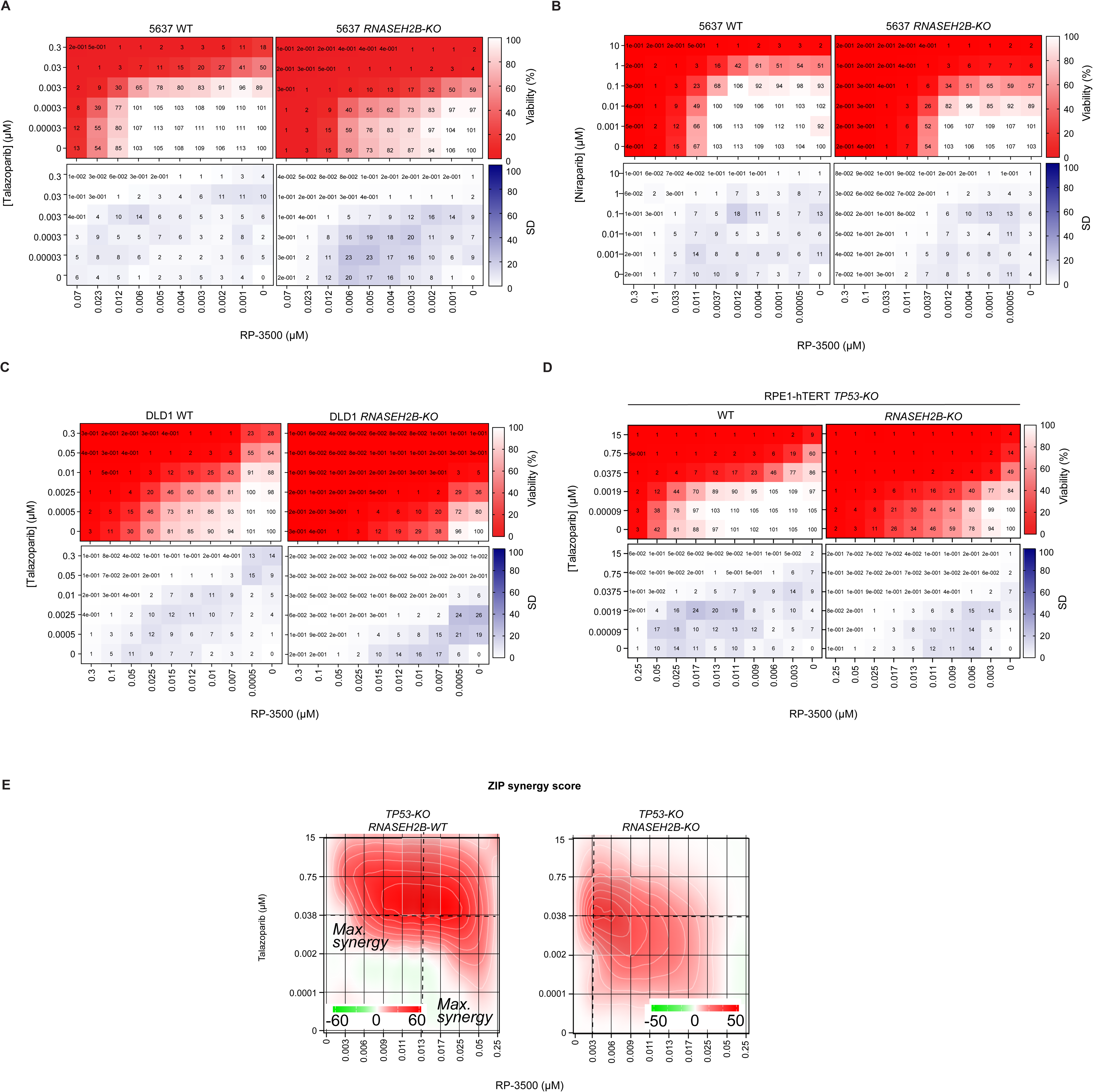
Related to Figure 2. Heat maps showing viability of 5637 (**A,B**), DLD1 (**C**) and RPE1 (**D**) WT and *RNASEH2B-KO* cells at indicated concentrations of RP-3500 (ATRi), talazoparib (PARPi, **A,C,D**), niraparib (PARPi, **B**), or their combinations. Top panels (white-to-red scale) visualize mean viability values from three independent biological replicates with corresponding SDs shown in bottom panels (white-to-purple scale). **E**. ZIP synergy scores in RPE1 *TP53-KO RNASEH2B-WT* and *TP53-KO RNASEH2B-KO* cells at various dose combinations of RP-3500 and talazoparib. Score ≥10 (red color) represents synergy, ≤-10 (green) antagonism. Dashed lines mark doses showing maximal synergy. Values were obtained by analyzing mean data from three independent biological replicates with SynergyFinder.

**Supplementary Figure 4.**
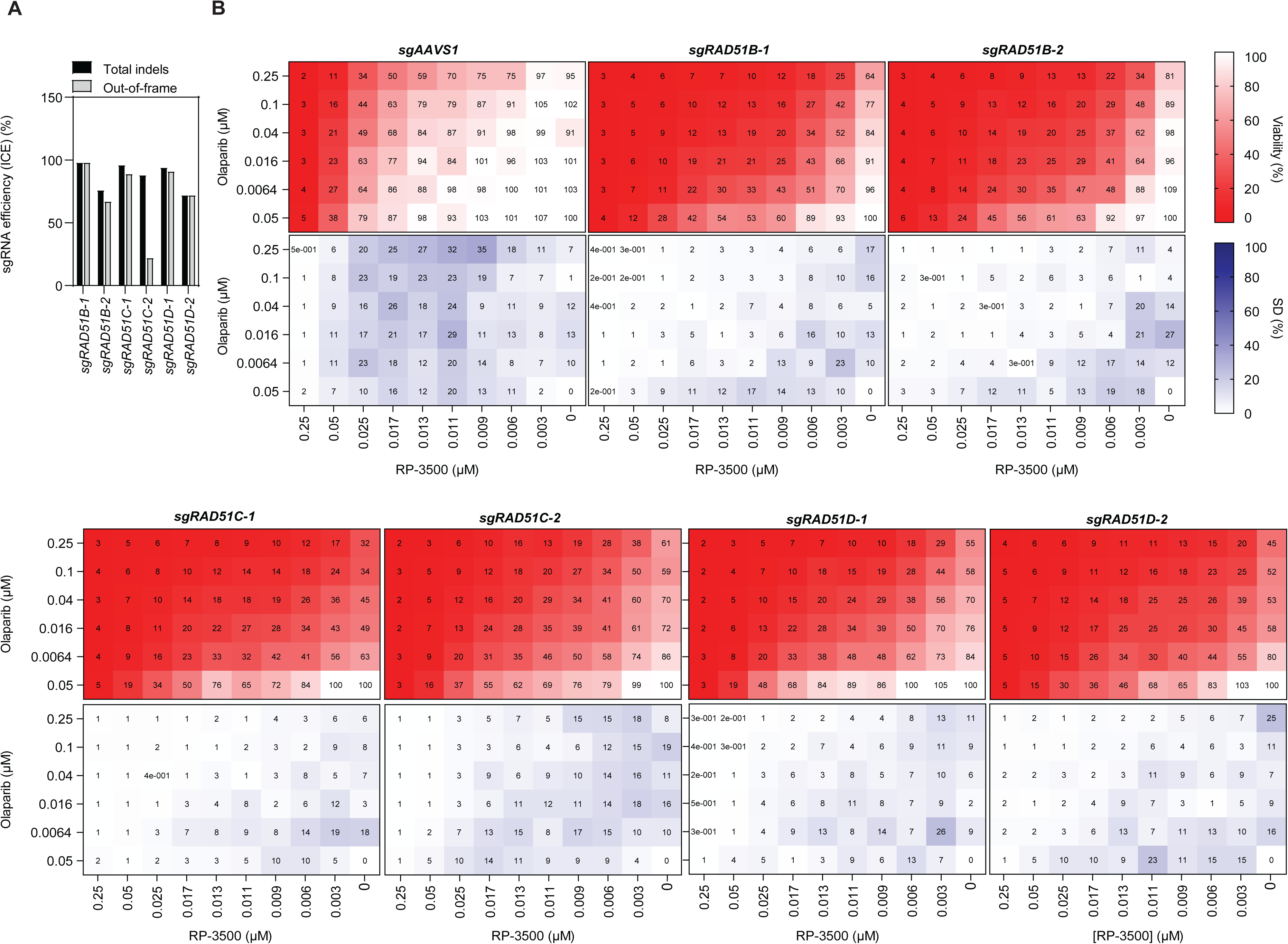
Related to Figure 3. **A.** CRISPR sgRNA editing efficiencies in experiments shown in Figure 3 as determined by ICE (Hsiau et al., 2018). The percentages of total indel mutations as well as only out-of-frame indels, which are more likely to induce a gene knockout, are shown. **B.** Heat maps showing viability of Cas9-expressing RPE1-hTERT TP53-KO cells transduced with the indicated sgRNAs treated with RP-3500 (ATRi), olaparib (PARPi) or their combination. Top panels (white-to-red scale) visualize mean viability values from three independent biological replicates with corresponding SDs shown in bottom panels (white-to-purple scale).

**Supplementary Figure 5.**
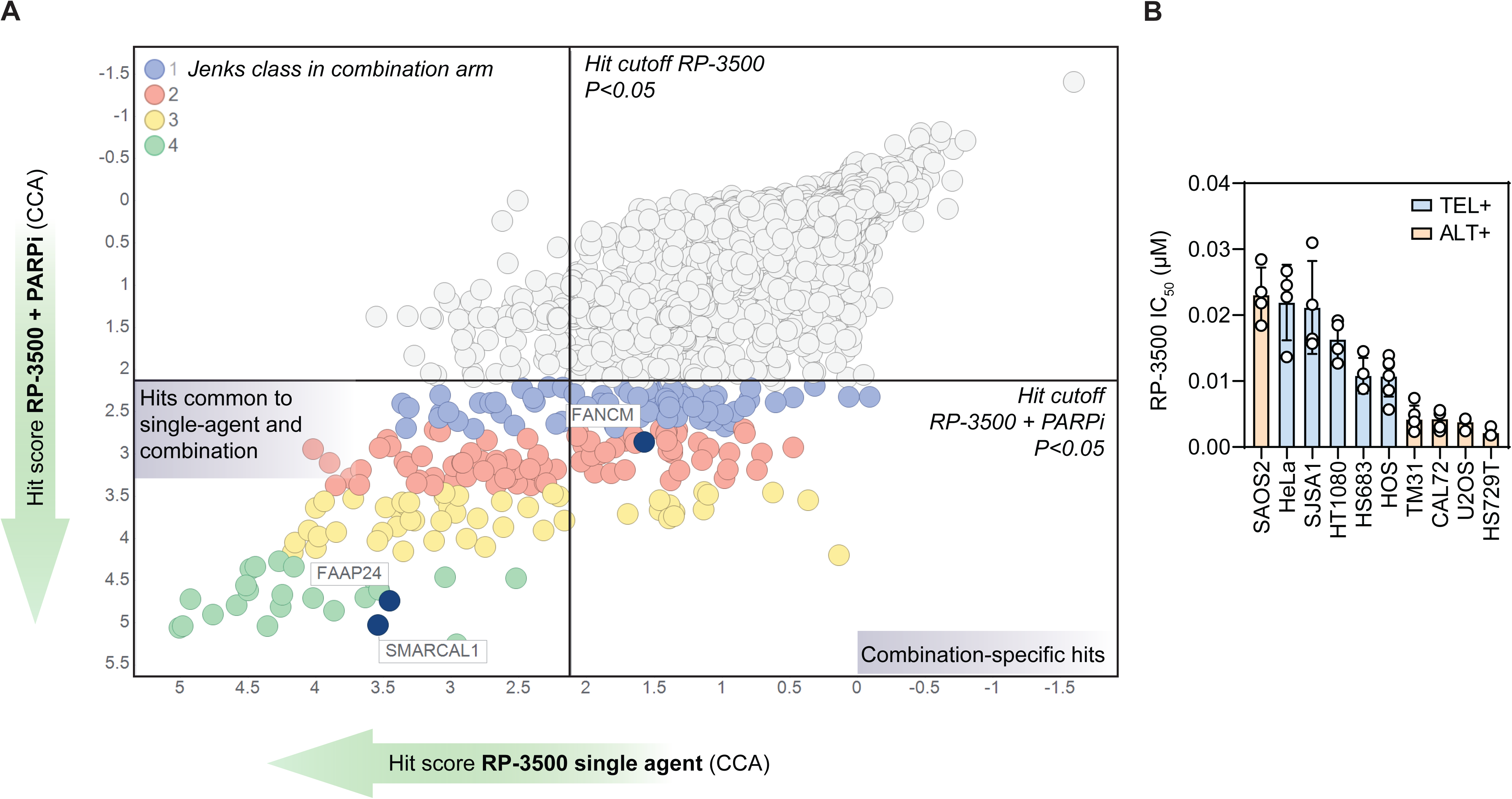
Related to Figure 4. **A.** ALT-related genes highlighted on the RP-3500 and RP-3500+PARPi screen diagram. **B.** Single agent RP-3500 sensitivity of a panel of ALT-positive (ALT+, orange bars) and telomerase-positive (TEL+, blue bars) cancer cell lines. Circles show RP-3500 IC_50_ from ≥3 independent biological replicates. Bars show mean±SD.

**Supplementary Figure 6.**
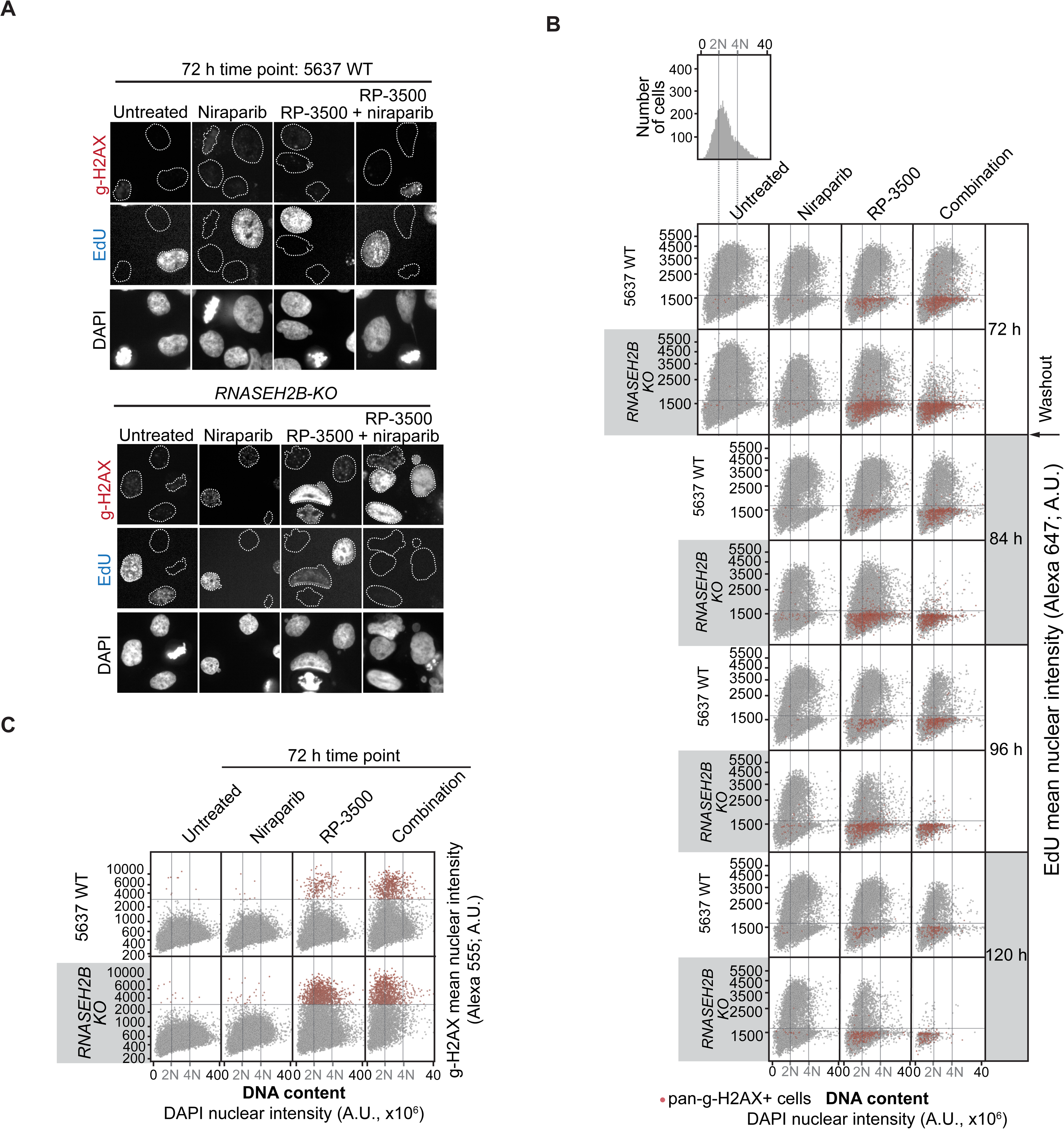
Related to Figure 5. **A.** Representative high-content microscopy images of WT and *RNASEH2B-KO* cells stained for γ-H2AX and EdU at the 72-hour timepoint for experiments shown in **B,C** and Figure 5A-C. DAPI – nuclear counterstain. **B.** Example cell cycle profiles for 5637 WT and *RNASEH2B-KO* cells from experiments shown in Figure 5A-C. For each cell, DNA content as measured by total DAPI nuclear intensity was plotted against the mean nuclear EdU intensity. Horizontal line in each panel indicates the cut-off for EdU+ cells and vertical lines show the approximate peak of 2N and 4N cell populations as determined from DNA content histograms (example shown in top-most panel). Pan-nuclear γ-H2AX cells are highlighted in red. **C.** An example of gating strategy to quantify pan-γ-H2AX+ cells in **B** and Figure 5B. DNA content was plotted against mean γ-H2AX nuclear intensity. Horizontal lines indicate the cut-off for γ-H2AX+ cells and vertical lines show the approximate peak of 2N and 4N cell populations as determined from DNA content histograms. All panels are representative of three independent biological replicates.

**Supplementary Figure 7.**
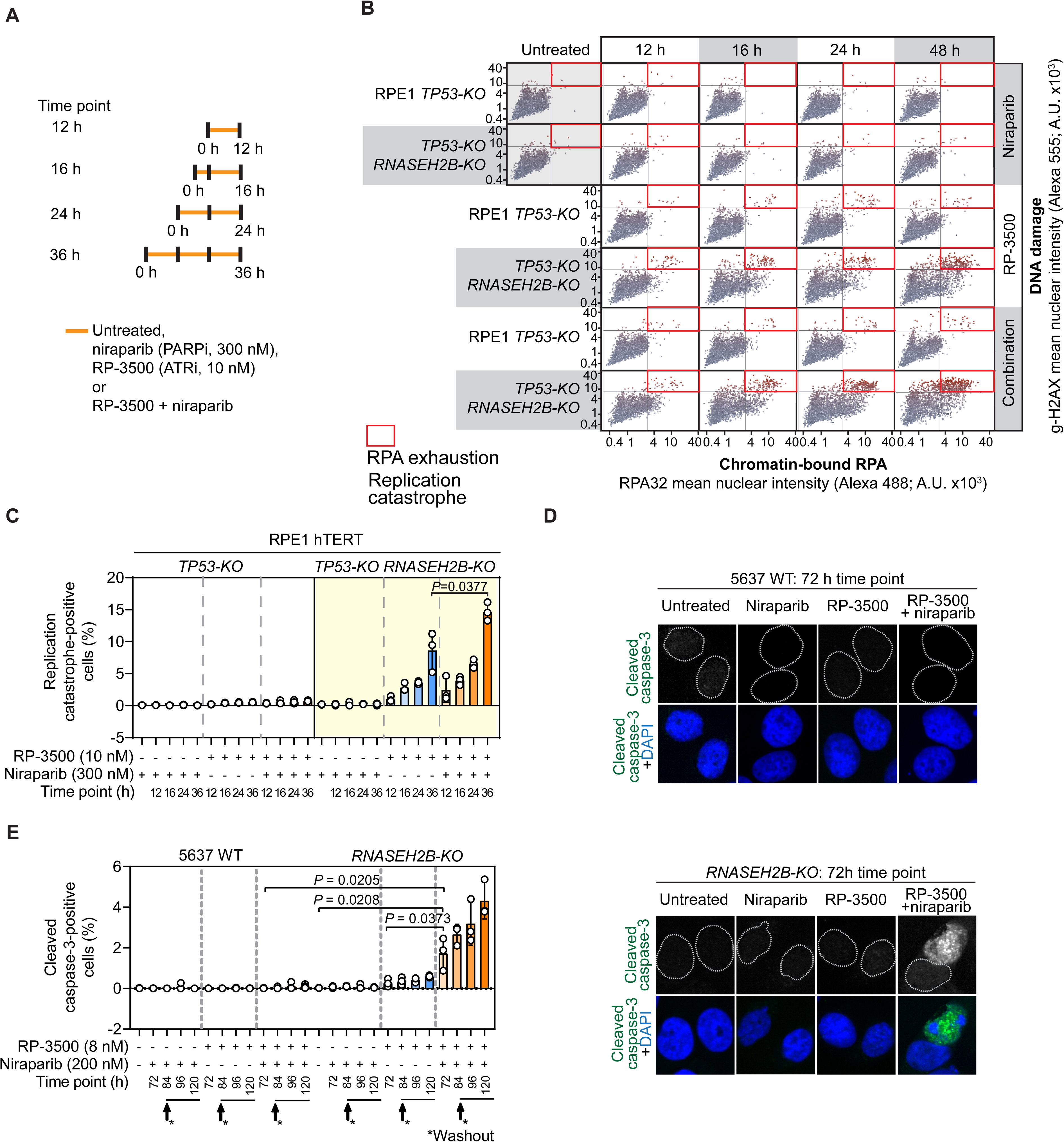
Related to Figure 5. **A.** Timeline of experiments shown in **B,C**. RPE1-hTERT *TP53-KO* or *TP53-KO RNASEH2B-KO* cells were subjected to indicated treatments for indicated amounts of time and processed for immunostaining. **B.** High-content microscopy quantification of immunofluorescence signals of chromatin-bound RPA32 and γ-H2AX in cells of indicated genotypes treated as shown in **A**. Each point represents a single nucleus, solid lines show cut-offs for γ-H2AX+ and RPA32+ cells. Replication catastrophe+ cells (RPA32+/γ-H2AX+) are outlined with red rectangles. Representative plots of three independent biological replicates. **C.** Percentages of replication catastrophe+ cells in the indicated samples. **D.** Representative (of three biological replicates) high-content microscopy images of 5637 WT and *RNASEH2B-KO* cells stained for cleaved caspase-3 (apoptotic cells; green); DAPI – nuclear counterstain. **E.** Automated high content microscopy quantification of 5637 WT and *RNASEH2B-KO* cells positive (+) for cleaved caspase-3 after indicated treatments and at indicated time points. Data in **C,E** are represented as follows: Circles are values from three independent biological replicates; Bars indicate mean ±SD. *P* value calculated with a two-tailed unpaired Student’s t-test.

**Supplementary Figure 8.**
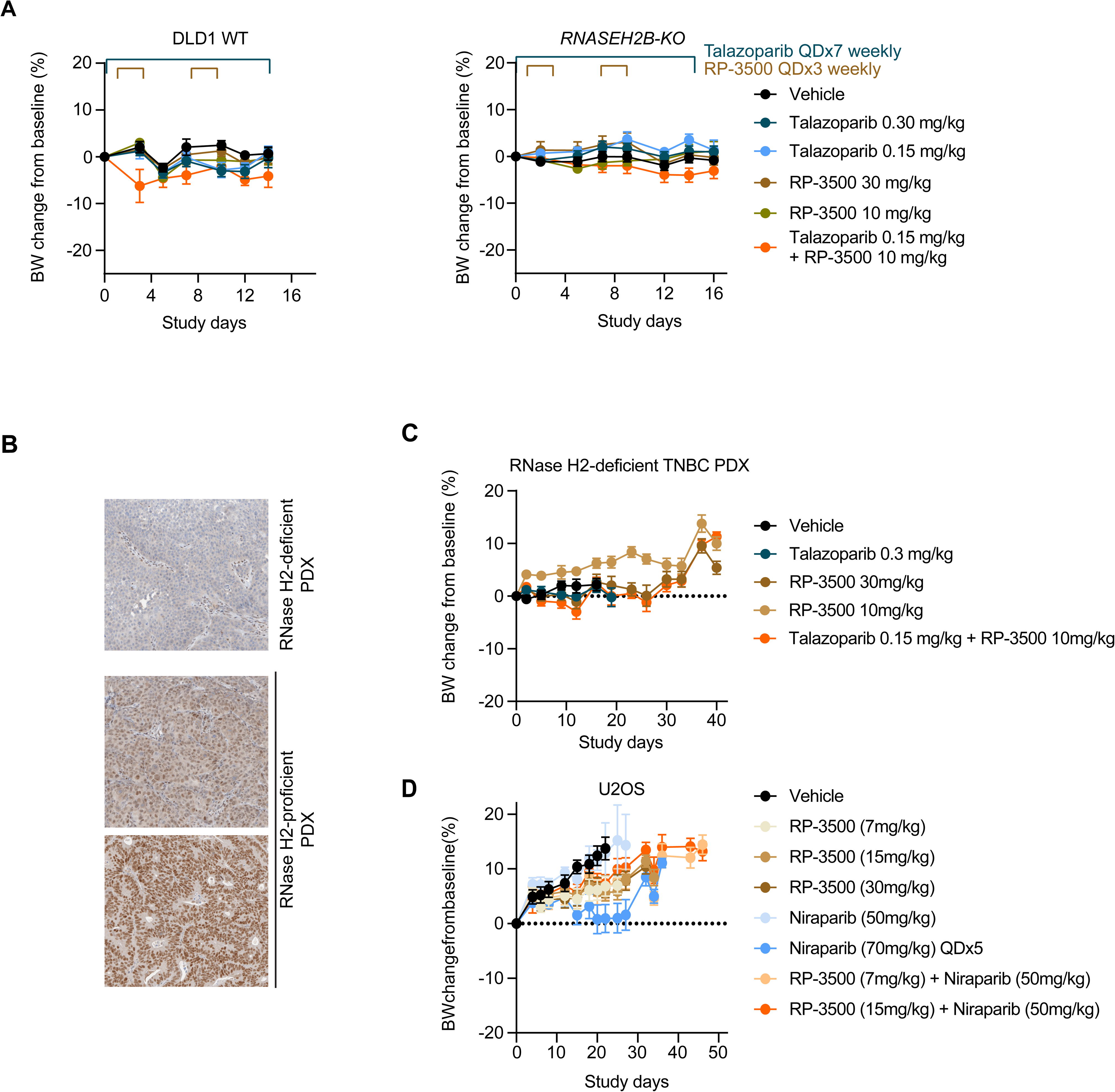
Related to Figure 6. **A.** Tolerability of RP-3500, talazoparib and the combination in DLD1 xenograft experiments in Figure 6A. Mean BW measures normalized to baseline are plotted for indicated cohorts. Error bars represent ±SEM, N=8 and 7 mice/group for WT and *RNASEH2B-KO*, respectively. **B.** Representative IHC images of RNase H2-deficient and -proficient PDX models. Deficient PDX on the top panel was used in the experiment shown in Figure 6B. **C.** Tolerability of RP-3500, talazoparib and the combination in the PDX experiment in Figure 6B. Mean BW measures normalized to baseline are plotted for indicated cohorts. Error bars represent ±SEM, N=9 mice/group. **D.** Tolerability of RP-3500, niraparib and the combination in the U2OS xenograft experiment in Figure 6C. Mean BW measures normalized to baseline are plotted for indicated cohorts. Error bars represent ±SEM, N=9 mice/group.

